# Multi-omics-based label-free metabolic flux inference reveals obesity-associated dysregulatory mechanisms in liver glucose metabolism

**DOI:** 10.1101/2021.06.21.449220

**Authors:** Saori Uematsu, Satoshi Ohno, Kaori Y. Tanaka, Atsushi Hatano, Toshiya Kokaji, Yuki Ito, Hiroyuki Kubota, Ken-ichi Hironaka, Yutaka Suzuki, Masaki Matsumoto, Keiichi I. Nakayama, Akiyoshi Hirayama, Tomoyoshi Soga, Shinya Kuroda

## Abstract

Glucose homeostasis is maintained by modulation of metabolic flux. Enzymes and metabolites regulate the involved metabolic pathways. Dysregulation of glucose homeostasis is a pathological event in obesity. Analyzing metabolic pathways and the mechanisms contributing to obesity-associated dysregulation *in vivo* is challenging. Here, we introduce OMELET: Omics-Based Metabolic Flux Estimation without Labeling for Extended Trans-omic Analysis. OMELET uses metabolomic, proteomic, and transcriptomic data to identify changes in metabolic flux, and to quantify contributions of metabolites, enzymes, and transcripts to the changes in metabolic flux. By evaluating the livers of fasting *ob*/*ob* mice, we found that increased metabolic flux through gluconeogenesis resulted primarily from increased transcripts, whereas that through the pyruvate cycle resulted from both increased transcripts and changes in substrates of metabolic enzymes. With OMELET, we identified mechanisms underlying the obesity-associated dysregulation of metabolic flux in liver.

**Highlights:**

- We created OMELET to infer metabolic flux and its regulation from multi-omic data.
- Gluconeogenic and pyruvate cycle fluxes increased in fasting *ob*/*ob* mice.
- Transcripts increases mediated the increase in gluconeogenic fluxes in *ob*/*ob* mice.
- Increases in transcripts and substrates enhanced pyruvate cycle flux in *ob*/*ob* mice.

## INTRODUCTION

Glucose homeostasis is tightly regulated to meet the energy requirements of vital organs and maintain health. Dysregulation of glucose homeostasis leads to metabolic diseases such as obesity and type 2 diabetes (Hotamisligil and Erbay, 2008; Kahn et al., 2006; Petersen et al., 2017). The liver plays a central role in glucose homeostasis by regulating various pathways of glucose metabolism, including gluconeogenesis and glycolysis (Han et al., 2016; Nordlie et al., 1999; Petersen et al., 2017). The liver is a major player in the pathophysiology of obesity (Charlton, 2004; Polyzos et al., 2019; Roden and Shulman, 2019). Fasting hyperglycemia in obesity is attributed to altered glucose metabolism in the liver due to insulin resistance. Because of the complex nature of the obesity-associated pathophysiology of glucose metabolism in liver, investigation of the dysregulation in this metabolic system requires data of multiple types that are obtained under the same condition.

Metabolic flux, the rate of turnover of molecules through a metabolic pathway, is a direct measure of the activity of the metabolic pathway (Jang et al., 2018). Metabolic flux is controlled by multiple regulators: enzymes, substrates, products, and cofactors. Enzymes are regulated by allosteric effectors and other factors such as post-translational modifications of enzymes. The amounts of enzymes are determined by the amounts of transcripts encoding the corresponding enzymes and other factors such as translation and protein degradation. To investigate metabolic flux and its complex regulation, the amounts of all the regulators of metabolic flux should be simultaneously measured because molecular interactions between metabolome layer and other multiple omic layers are mutually connected (Wiley, 2011; Yugi et al., 2014). The amounts of enzymes can be measured by mass spectrometry-based proteomics, transcripts for enzymes by RNA sequencing, and the amounts of substrates, products, cofactors, and allosteric effectors by mass spectrometry-based metabolomics. We developed a method of trans-omic analysis based on direct molecular interactions to construct a multilayered biochemical network using simultaneously obtained multi-omic data (Egami et al., 2021; Kawata et al., 2018; Kokaji et al., 2020; Yugi et al., 2014, 2016). This trans-omic approach was not used to infer metabolic fluxes.

The standard method for measuring metabolic flux is isotopic labeling, in which isotopic tracers are introduced into cells or living animals (Hasenour et al., 2015, 2020; Hiller and Metallo, 2013; Quek et al., 2010). To analyze the regulation of metabolic flux in a non-steady state, we developed a kinetic trans-omic analysis that uses data from isotopic labeling experiments and inferred metabolic flux together with contributions of regulators to changes in metabolic flux across a multi-layered network (Ohno et al., 2020).

A problem with the application of isotope labeling experiments in living animals is that the addition of isotopic tracers can perturb the relevant metabolic flux, resulting in different metabolic states (Previs and Kelley, 2015). For example, when ^13^C-propionate or ^13^C-lactate is used as an isotopic tracer for measurement of gluconeogenic flux, the measured metabolic fluxes differ between tracers because the administration of ^13^C-propionate increases metabolic flux through the pyruvate cycle (Perry et al., 2016). Therefore, the addition of isotopic tracers inevitably results in experimentally induced changes in metabolic flux. To avoid the effects of the addition of isotopic tracers, a method to infer metabolic flux without using isotopic tracers is needed to be developed.

Here, we present a method that we termed as Omics-Based Metabolic Flux Estimation without Labeling for Extended Trans-omic Analysis (OMELET). OMELET infers metabolic fluxes in each condition from metabolomic, proteomic, and transcriptomic data, which are simultaneously obtained from the same individual samples, identifies changes in metabolic flux between the conditions, and (iii) quantifies contributions of regulators to the changes in metabolic flux. We obtained metabolomic, proteomic, and transcriptomic data from the livers of wild-type (WT) and leptin-deficient obese (*ob*/*ob*) mice in the fasting state and four hours after oral glucose administration. By applying OMELET to the experimental data, we inferred metabolic fluxes in each condition, and quantified contributions of regulators to changes in metabolic flux between the conditions. In the fasting state, metabolic fluxes through reactions in gluconeogenesis and the pyruvate cycle increased in *ob*/*ob* mice compared to WT mice. The increased metabolic fluxes through reactions in gluconeogenesis were caused by increased transcripts. In contrast, in the pyruvate cycle, the increased metabolic fluxes through pyruvate kinase (PK) involved increased transcripts and that through phosphoenolpyruvate carboxykinase (PEPCK) was caused by increased substrates. We also quantified the contributions of regulators to changes in metabolic flux resulting from oral glucose administration. In response to oral glucose administration, although the metabolic flux through PK did not change in both WT and *ob*/*ob* mice, the regulation of metabolic flux changed: PK flux was regulated by increased ATP as an allosteric inhibitor in WT mice, and by decreased PK-encoding transcript in *ob*/*ob* mice. Thus, OMELET provided quantitative mechanistic insights into obesity-associated differences in metabolic regulation in liver without using isotopic tracers.

## RESULTS

### Overview of the application of OMELET to study glucose metabolism

In this study, we developed OMELET to infer metabolic flux using multi-omic data without using isotopic tracers, identify changes in metabolic flux between conditions, and quantify contributions of regulators to the changes in metabolic flux (Figure 1). We applied this method to evaluate the differences in metabolic flux in liver between WT and *ob*/*ob* mice, and the dysregulatory mechanisms associated with obesity. Additionally, we evaluated the differences in metabolic flux between the fasting state and 4-hours after oral glucose administration for both WT mice and *ob*/*ob* mice. We had four conditions: WT in the fasting state, WT after oral glucose administration, *ob*/*ob* in the fasting state, and *ob*/*ob* after oral glucose administration. In each condition, we measured the amounts of metabolites, enzymes, and transcripts in liver samples from each mouse. We orally administered glucose to mice that had fasted for 16 hours and collected livers before and four hours after oral glucose administration (Figure 1: Experimental data).

**Figure 1.**
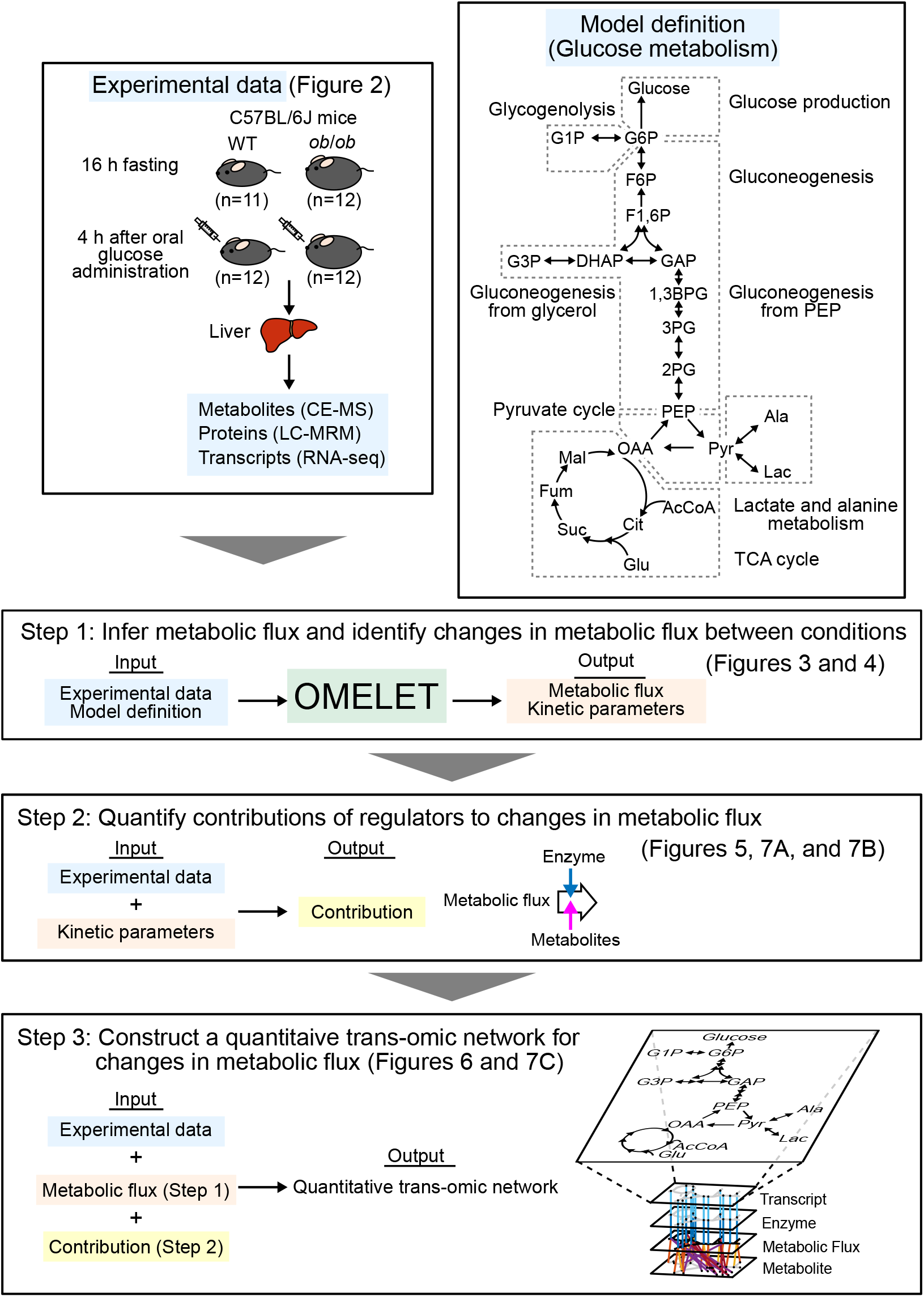
Overview of the application of OMELET to study glucose metabolism. (Top right) Overview of glucose metabolism (see Table S1 for definitions of metabolites). (Top left) Experimental data are acquired from livers of WT and *ob*/*ob* mice under fasting conditions and after oral glucose administration. These data serve as the input for Step 1.

By applying OMELET to the experimental data, we inferred metabolic fluxes in the glucose metabolism in each condition, and identified changes in metabolic flux between the conditions (Figure 1: Step 1). We focused on reactions in the glucose metabolism and inferred metabolic fluxes through the reactions in glycogenolysis, gluconeogenesis, lactate and alanine metabolism, the pyruvate cycle, and the TCA cycle (Figure 1; Table S1).

The changes in metabolic fluxes between conditions are caused by the changes of regulators such as enzymes and metabolites. To investigate which regulators caused changes in metabolic flux between the conditions, we quantified the contributions of the regulators to changes in metabolic flux from experimental data and kinetic parameters obtained in Step 1 (Figure 1: Step 2).

By integrating changes in the experimental data, changes in metabolic flux (Figure 1: Step 1), and contributions of the regulators to the changes in metabolic flux between the conditions (Figure 1: Step 2), we constructed a quantitative trans-omic network of the glucose metabolism in liver, which represents changes in metabolic flux and the regulation across multi-omic layers associated with obesity (Figure 1: Step 3).

### Metabolomic, proteomic, and transcriptomic analysis of glucose metabolism in livers from WT and *ob*/*ob* mice in the fasting state and after oral glucose administration

We obtained metabolomic, proteomic, and transcriptomic data from livers of WT and *ob*/*ob* mice in the fasting state and four hours after oral glucose administration (Figure 2). The dynamics of blood glucose and insulin concentrations differed between WT and *ob*/*ob* mice, consistent with obesity phenotype of the *ob*/*ob* mice (Figures S1A and S1B). However, both groups reached a steady state four hours after oral glucose administration. The transcriptomic data were reported in our previous studies (Egami et al., 2021; Kokaji et al., 2020), and the metabolomic and proteomic data were newly obtained in this study (Materials and Methods). We selected 28 metabolites, 15 enzymes, and 17 transcripts relevant to glucose metabolism from the metabolomic, proteomic, and transcriptomic data, respectively (Table S2). We defined transcript, enzyme, and reaction names as follows; transcript names are italicized with only the first letter in upper-case (*e.g.*, *Pklr*), enzyme names are not italicized with only the first letter in upper-case (*e.g.*, Pklr), and reaction names are not italicized with all letters in upper-case (*e.g.*, PK). Principal component analysis of the metabolites, enzymes, and transcripts showed that the first principal components captured differences between WT and *ob*/*ob* mice, and the second principal components captured changes by oral glucose administration (Figure S1C). The principal component analysis indicated that the differences between the genotypes, represented by principal component 1, exceeded the differences within the genotypes related to oral glucose administration, represented by principal component 2. Indeed, the principal component 1 represented ≥50% of the variance (58% for metabolites, 81% for enzymes, and 50% for transcripts) and principal component 2 represented ≤15% of the variance (11% for metabolites, 8% for enzymes, and 15% for transcripts).

**Figure 2.**
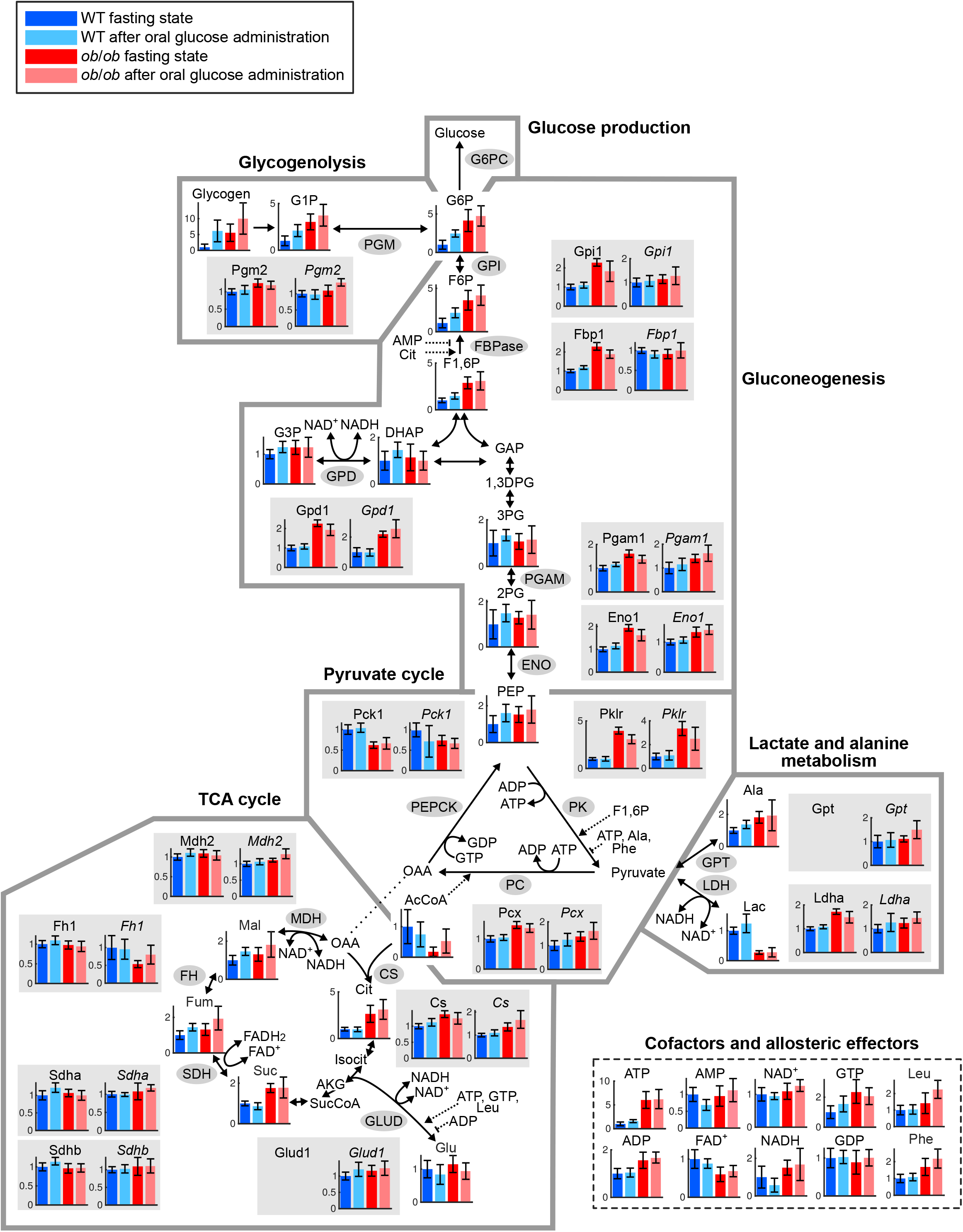
Metabolomic, proteomic, and transcriptomic analysis of glucose metabolism in livers from WT and *ob*/*ob* mice in the fasting state and after oral glucose administration. Measured molecules (metabolites, enzymes, and transcripts) mapped onto the glucose metabolism in liver. Irreversible reactions are shown with one-headed arrows; reversible reactions are shown with double-headed arrows. Allosteric activation and inhibition are shown with dotted one-headed and dotted bar-headed arrows, respectively. The bars and error bars in each molecule represent the mean ± SD normalized to the mean of the data from WT in the fasting state. Enzymes and transcript results are shaded in gray. G6pc was not measured at the protein or transcript level; Gpt and Glud1 were not measured at the protein level. Definitions of the metabolites, enzymes, and transcripts are described in Table S2.

We compared amounts of molecules between the conditions and defined increased and decreased molecules between the conditions. Molecules that showed an FDR-adjusted p value (q value) less than 0.05 were defined as significantly changed molecules. Among them, molecules that showed a fold change larger than 1.5 and smaller than 0.67 between the conditions were defined as increased and decreased molecules, respectively (Tables S2 and S3).

Consistent with the greatest separation between the genotypes by principal component analyses, we observed the greatest number of molecules in glucose metabolism differed between WT and *ob*/*ob* mice (Tables S2 and S3). Comparing WT and *ob*/*ob* mouse livers in the fasting state showed that increased metabolites and enzymes in *ob*/*ob* mice included those in glycogenolysis and gluconeogenesis. After oral glucose administration, differences in metabolites in *ob*/*ob* mouse livers compared to WT mouse livers partially overlapped with the differences between the genotypes in the fasting state, however, only three increased enzymes were observed. WT mouse livers showed increases in metabolites of glycogenolysis and gluconeogenesis following glucose administration. In *ob*/*ob* mouse livers, no metabolites in glucose metabolism were significantly changed by glucose administration. Neither WT or *ob*/*ob* mice had any changes in enzymes or transcripts when livers from fasting mice were compared to livers from mice of the same genotype after oral glucose administration.

The amounts of metabolites, enzymes, and transcripts do not directly reflect metabolic flux and its regulation; however, these data contain sufficient information to infer metabolic flux and its regulation. Therefore, we developed a method to infer metabolic flux and its regulation using the metabolomic, proteomic, and transcriptomic data.

### Inference of metabolic fluxes by OMELET

OMELET is a probability-based model that incorporates metabolomic, proteomic, and transcriptomic data and uses kinetic equations to predict the parameters of metabolic flux, the elasticity coefficients, and the protein turnover coefficients for each reaction (Figure 3). The advantages of OMELET are that metabolic flux can be inferred without using isotopic tracers, and that the regulation of metabolic flux can be determined from the kinetic parameters inferred by OMELET. The inputs of OMELET are the experimental data of the amounts of metabolites ***x***, enzymes ***e***, and transcripts ***t*** from the same mouse in each condition as well as model definitions, which are a stoichiometric matrix of the target metabolic pathway and information on cofactors and allosteric effectors for each reaction. The outputs are metabolic fluxes ***v*** in the target metabolic pathway in each condition, elasticity coefficients ***ϵ*** , and protein turnover coefficients ***β*** . The elasticity coefficient is the change in metabolic flux in response to infinitesimal changes in metabolites normalized to a reference condition. OMELET is based on a Bayesian method that calculates posterior probability of the output parameters *p*(***v***, ***ϵ***, ***β***|***x***, ***e***, ***t***) by updating prior probability of parameters [*p*(***v***|***u***), *p*(***ϵ***), and *p*(***β***)] and the hyperprior of independent flux *p*(***u***|***μ***^***u***^) . The posterior probability of the output parameters is achieved by evaluating likelihoods *p*(***e***, ***t***|***x***, ***v***, ***β***, ***ϵ***) of the proteomic and transcriptomic data under a given metabolomic data and parameter set including metabolic flux (Materials and Methods). Metabolic flux ***v*** in a given metabolic pathway under a steady-state condition can be written as a linear combination of independent flux ***u***, and the prior probability of metabolic flux *p*(***v***|***u***) is assumed to follow a multivariate normal distribution 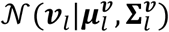 We used the elasticity coefficients ***ϵ*** and protein turnover coefficients ***β*** to calculate the contributions of regulators to the changes in metabolic flux between the conditions. Thus, OMELET enabled identification of the specific reaction with changes in metabolic flux between the conditions and the extent to which specific regulators, such as changes in the amounts of enzymes and metabolites, contributed the inferred differences in metabolic flux between the conditions.

**Figure 3.**
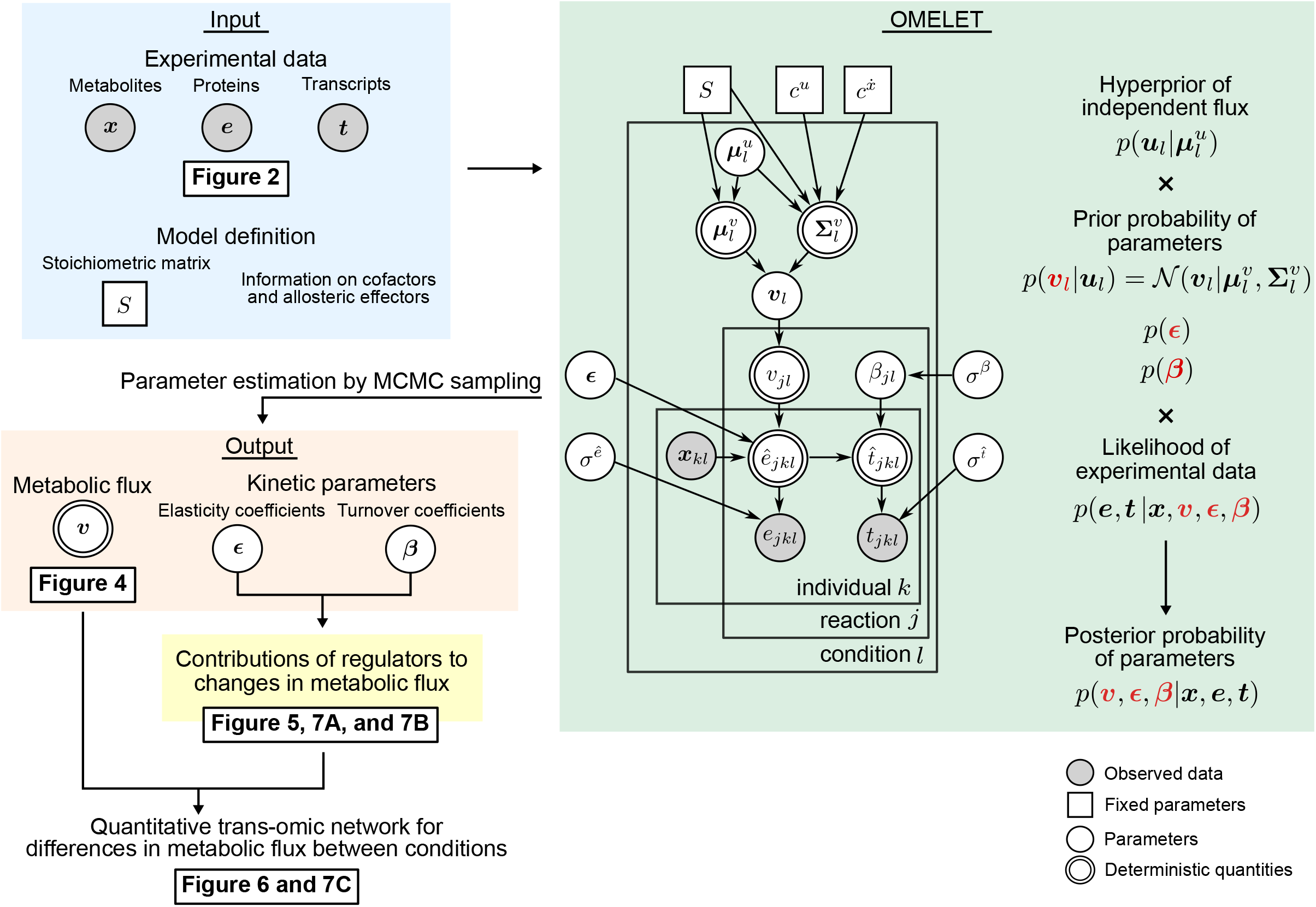
Inference of metabolic fluxes by OMELET. Overview of the workflow for the inference of metabolic flux. Shaded rectangles represent the inputs (light blue), OMELET (light green), the outputs (orange), and contributions (yellow). The inputs of OMELET were the experimental data of the amounts of metabolites ***x***, enzymes ***e***, and transcripts ***t*** from the same individual mice in each condition as well as model definitions including stoichiometric matrix and information on cofactors and allosteric effectors. The outputs were metabolic fluxes in the glucose metabolism ***v*** in each condition, elasticity coefficients ***ϵ***, and protein turnover coefficients ***β***. The output parameters are colored in red. In the graphical model, the plate indicates that the group-level structure holds for all the analyzed reactions *j* ∈ *R*, samples *k* = 1, … , *n*_*l*_, and conditions *l* = 1. … , *g*. The arrows denote conditional dependences between two nodes representing the generating processes (Materials and Methods). Shaded circles, unshaded squares, single-bordered circles, and double-bordered circles represent observed data, fixed parameters, parameters, and deterministic quantities, respectively. Using the kinetic parameters including elasticity coefficients and turnover coefficients, we can calculate contributions of regulators to changes in metabolic flux between conditions. See also Materials and Methods.

We validated the performance of OMELET by applying it to simulated datasets of the amounts of metabolites, enzymes, and metabolic fluxes in five conditions from a kinetic model representing yeast glycolysis (Messiha et al., 2014; Smallbone et al., 2013) (Figure S2; Table S4). The metabolic fluxes inferred by OMELET highly correlated with those generated by steady-state simulations of the kinetic model for different yeast mutants (Figure S2), indicating that OMELET accurately identified the difference in metabolic fluxes across the reactions in glycolysis in each condition and the changes in metabolic flux among the mutants.

### Inference of metabolic fluxes in the glucose metabolism in liver of WT and *ob*/*ob* mice in the fasting state and after oral glucose administration

Because the blood glucose and insulin were constant both in WT and *ob*/*ob* mice in the fasting state and four hours after oral glucose administration (Figures S1A and S1B), we assumed steady-state conditions for glucose metabolism in livers of WT and *ob*/*ob* mice. By applying OMELET to the experimental data, we inferred metabolic fluxes in glucose metabolism in four conditions: WT and *ob*/*ob* mice in the fasting state and after oral glucose administration (Figures 4A and S3; Table S5). The posterior distributions of the metabolic fluxes were obtained by fitting the model to the experimental data for the amounts of enzymes and transcripts in each condition (Figure S4). We assumed that liver produces glucose through gluconeogenesis, but not consume glucose through glycolysis, in all the conditions analyzed according to the previous studies (Jin et al., 2013; Turner et al., 2005). We fixed the direction of the reaction from glucose 6-phosphate (G6P) to glucose, mediated by glucose-6-phosphatase and indicated in the model as G6PC for glucose production. A metabolic flux through each reaction was simultaneously inferred in all the conditions as the relative value to that through G6PC in WT mice in the fasting state (Materials and Methods).

**Figure 4.**
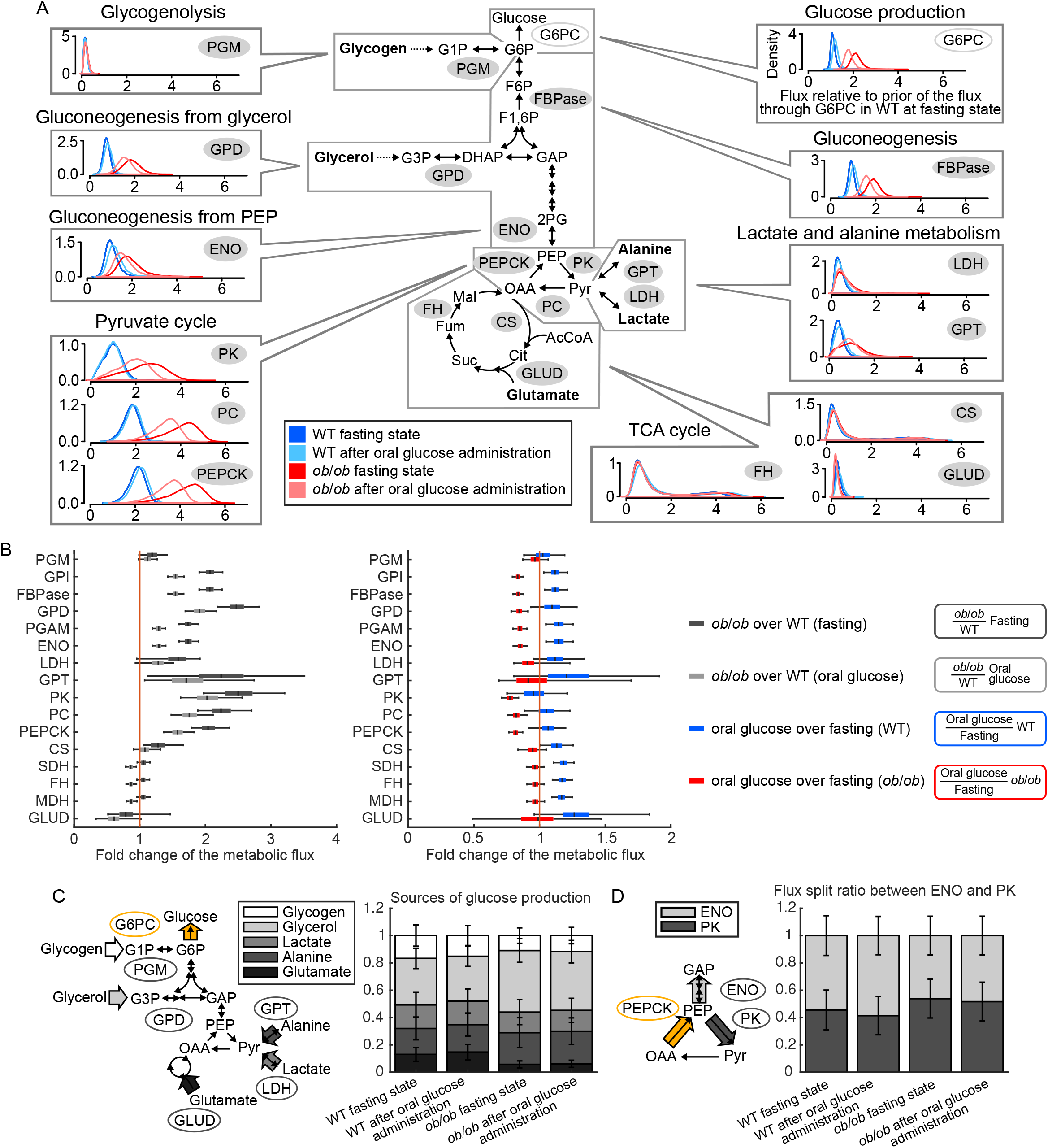
Inference of metabolic fluxes in the glucose metabolism in liver of WT and *ob*/*ob* mice in the fasting state and after oral glucose administration. (A) Posterior distributions of the metabolic fluxes in the glucose metabolism. Each box contains four density plots corresponding to four different conditions. Metabolic flux through each reaction is inferred relative to the mean of the prior for the metabolic flux through G6PC in WT mice in the fasting state. Only representative reactions (shaded gray circles in the map) in each pathway are presented See Figure S3 and Table S5 for complete reaction data. (B) Fold changes of the metabolic flux of *ob*/*ob* mice over that of WT mice in the fasting state (black bars) and after oral glucose administration (gray bars) in each reaction, and fold changes of the metabolic flux after oral glucose administration over that in the fasting state in WT mice (blue bars) and *ob*/*ob* mice (red bars) in each reaction. The median of the posterior distribution from OMELET is represented by a black line within the box for each reaction, the box extends from the lower to the 25^th^ and 75^th^ percentiles, and the whiskers extend to 2.5^th^ and 97.5^th^ percentiles to cover 95% of the data. The vertical orange line indicates the boundary where a fold change equals one. (C) Sources of glucose production. The stacked bars and error bars represent the mean ± SD of the proportions of glycogen, glycerol, lactate, alanine, and glutamate to the glucose production. The proportions of the sources are calculated from the proportion of metabolic fluxes through PGM, GPD, LDH, GPT, and GLUD, respectively (black circle) to that through G6PC (yellow circle). (D) Flux split ratios between ENO and PK reactions. The stacked bars and error bars represent the mean ± SD of the proportions of the metabolic fluxes through ENO and PK to that through PEPCK.

The metabolic flux through G6PC as glucose production is the sum of that through phosphoglucomutase (PGM) in glycogenolysis and fructose-1,6-bisphosphatase (FBPase) in gluconeogenesis. In WT mice in the fasting state, the 95% credible interval of the posterior distribution of the metabolic flux through FBPase (0.67 to 1.3) was not overlapped with that through PGM (0.05 to 0.38) (Figures 4A and S3; Table S5), suggesting that glucose production depended on gluconeogenesis. The small metabolic flux through PGM was consistent with the depletion of glycogen in WT mice in the fasting state (Figure 2). The metabolic flux through FBPase was further divided into that through glycerol-3-phosphate dehydrogenase (GPD) and alpha-enolase (ENO), which represented the metabolic flux through gluconeogenesis from glycerol and phosphoenolpyruvate (PEP), respectively. The metabolic flux through ENO (median of the posterior distribution: 1.1) and GPD (median: 0.75) were not significantly different (Figures 4A and S3; Table S5), indicating that both PEP and glycerol are equally used for glucose production. In the pyruvate cycle, the metabolic flux through PEPCK (median: 2.1) was larger than that through ENO (median: 1.1), and those through pyruvate carboxylase (PC) (median: 1.8) and PK (median: 0.98) were not significantly different from that through ENO (median: 1.1) (Figures 4A and S3; Table S5). This result suggested that PEP was equally used for glucose production and return to pyruvate. The metabolic flux through PK (median: 0.98) was not significantly different from those through alanine aminotransferase (GPT) (median: 0.41) and lactate dehydrogenase (LDH) (median: 0.35), suggesting that pyruvate synthesis is equally contributed by the influxes from PEP, alanine, and lactate through PK, GPT, and LDH, respectively. In the TCA cycle, 95% credible intervals of the metabolic fluxes were large (0.21 to 4.1 on average) compared to those through other reactions in glucose metabolism (0.54 to 0.15 on average), indicating that the metabolic fluxes through reactions in the TCA cycle were not reliably determined. The metabolic fluxes in glucose metabolism in the liver of fasting WT mice inferred by OMELET were consistent with those by the previous metabolic flux analyses of fasting WT mice except for those in the TCA cycle (Burgess et al., 2015; Hasenour et al., 2015, 2020; Satapati et al., 2012; Wang et al., 2020) (Figures S5A and S5B).

For *ob*/*ob* mice in the fasting state, we calculated fold changes of the metabolic flux of *ob*/*ob* mice over that of WT mice for each reaction (Figure 4B, black bars). Although the TCA cycle fluxes were not reliably determined in the individual conditions including fasting WT and *ob*/*ob* mice (Figure 4A), the fold changes of the metabolic flux of fasting *ob*/*ob* mice over that of fasting WT mice were inferred with relatively narrow 95% credible intervals (0.89 to 1.3 average) (Figure 4B; Table S5). The fold changes of metabolic fluxes through reactions in gluconeogenesis (median: 2.0 on average) and the pyruvate cycle (median: 2.3 on average) were larger than those in glycogenolysis (median: 1.2 on average) and the TCA cycle (median: 1.1 on average) (Figure 4B; Table S5). The metabolic flux through G6PC, glucose production, is a sum of the metabolic flux through PGM, GPD, LDH, GPT, and glutamate dehydrogenase (GLUD) multiplied by the number of carbon atoms of the substrates. We quantified the fraction of sources of the glucose production by calculating the proportion of the metabolic flux through PGM, GPD, LDH, GPT, and GLUD to that through G6PC, which represent glycogen, glycerol, lactate, alanine, and glutamate as sources, respectively (Figure 4C; Table S5). The median of the fraction of glucose production from glycerol was 34% in fasting WT mice and 45% in fasting *ob*/*ob* mice (Figure 4C; Table S5), and the 95% credible interval of the distribution of the fraction of fasting *ob*/*ob* mice minus that of fasting WT mice (5.6% to 17%) was greater than zero (Figure S6A; Table S5). This result suggested that fat accumulation in the liver of *ob*/*ob* mice increased the supply of glycerol as a precursor for glucose. The median of the fraction of glucose production from alanine was 19% in fasting WT mice and 24% in fasting *ob*/*ob* mice, whereas that from lactate was 16% in WT and 13% in *ob*/*ob* mice (Figures 4C and S6A; Table S5), implying that the contributions of alanine and lactate to glucose production did not change in *ob*/*ob* mice. The median of the fraction of glucose production from glycogen was 15% in WT and 10% in *ob*/*ob* mice, whereas that from glutamate was 12% in WT mice and 5.3% in *ob*/*ob* mice. Collectively, the fraction of glucose production from glycerol increased in *ob*/*ob* mice, whereas that from glycogen and glutamate decreased. To evaluate the efficiency of using carbons of PEP for glucose production through gluconeogenesis rather than for pyruvate through the pyruvate cycle, we calculated the flux split ratios between ENO and PK reactions (Figure 4D; Table S5) and compared them between fasting WT and *ob*/*ob* mice (Figure S6B; Table S5). Compared to WT mice, the ratio of the metabolic flux through PK in *ob*/*ob* mice was higher and was similar to that through ENO, indicating less efficient use of PEP as a source of glucose in *ob*/*ob* mice than in WT mice. Because the pyruvate cycle including PK is known as a futile cycle, in which no net PEP accumulation occurs but energy is used, the increased ratio of PK flux over ENO flux in *ob*/*ob* mice is likely to cause a futile ATP dissipation. We compared the fold changes of metabolic fluxes through reactions in the glucose metabolism of *ob*/*ob* mice over that of WT mice inferred by OMELET with those in the previous metabolic flux analyses in fasting *ob*/*ob* mice (Turner et al., 2005) and high-fat diet-induced obese mice (Patterson et al., 2016; Satapati et al., 2012) (Figures S5C and S5D). With OMELET, we found a larger increase in gluconeogenic flux than that in the previous studies (Satapati et al., 2012; Turner et al., 2005), and smaller increase in glycogenolysis flux than those in the previous studies (Satapati et al., 2012; Turner et al., 2005). Other differences in glucose metabolism between WT and *ob*/*ob* mice were consistent among the four studies.

To evaluate the effect of oral glucose administration on glucose metabolism, we calculated fold changes of the metabolic fluxes after oral glucose administration over those in the fasting state in WT mice (Figure 4B, blue bars) and *ob*/*ob* mice (Figures 4B, red bars). Orally administered glucose triggered a slight increase in the metabolic fluxes through most reactions in WT mice and a decrease in *ob*/*ob* mice. An exception was the metabolic flux through PK, which did not significantly change in WT mice and decreased slightly in *ob*/*ob* mice. Neither WT nor *ob*/*ob* mice exhibited much change from the fasting state in the sources of glucose production or the flux split ratio between ENO and PK in response to oral glucose administration (Figures 4C, 4D, and S6). These results suggested the differences in the metabolic flux between WT and *ob*/*ob* mice in the fasting state were maintained after oral glucose administration. However, the effect of oral glucose on the metabolic flux was opposite within each genotype: The metabolic fluxes slightly increased in WT mice and decreased in *ob*/*ob* mice by oral glucose administration.

### Contributions of regulators to changes in metabolic flux between fasting WT and *ob/ob* mice

Flux through reactions involved in metabolism is controlled by the enzymes, substrates, products, cofactors, and allosteric effectors. Each of these can be considered a “regulator” of the reaction. We quantified contributions of the regulators to changes in metabolic flux between the conditions (Figures 5 and S7; Table S6). The concept of the contribution is to partition the cause of changes in metabolic flux between conditions into underlying changes in the amounts of regulators. The contribution was calculated based on propagation of uncertainty of regulators’ amounts to metabolic flux, and a similar approach was described in a previous study (Hackett et al., 2016) (Materials and Methods).

**Figure 5.**
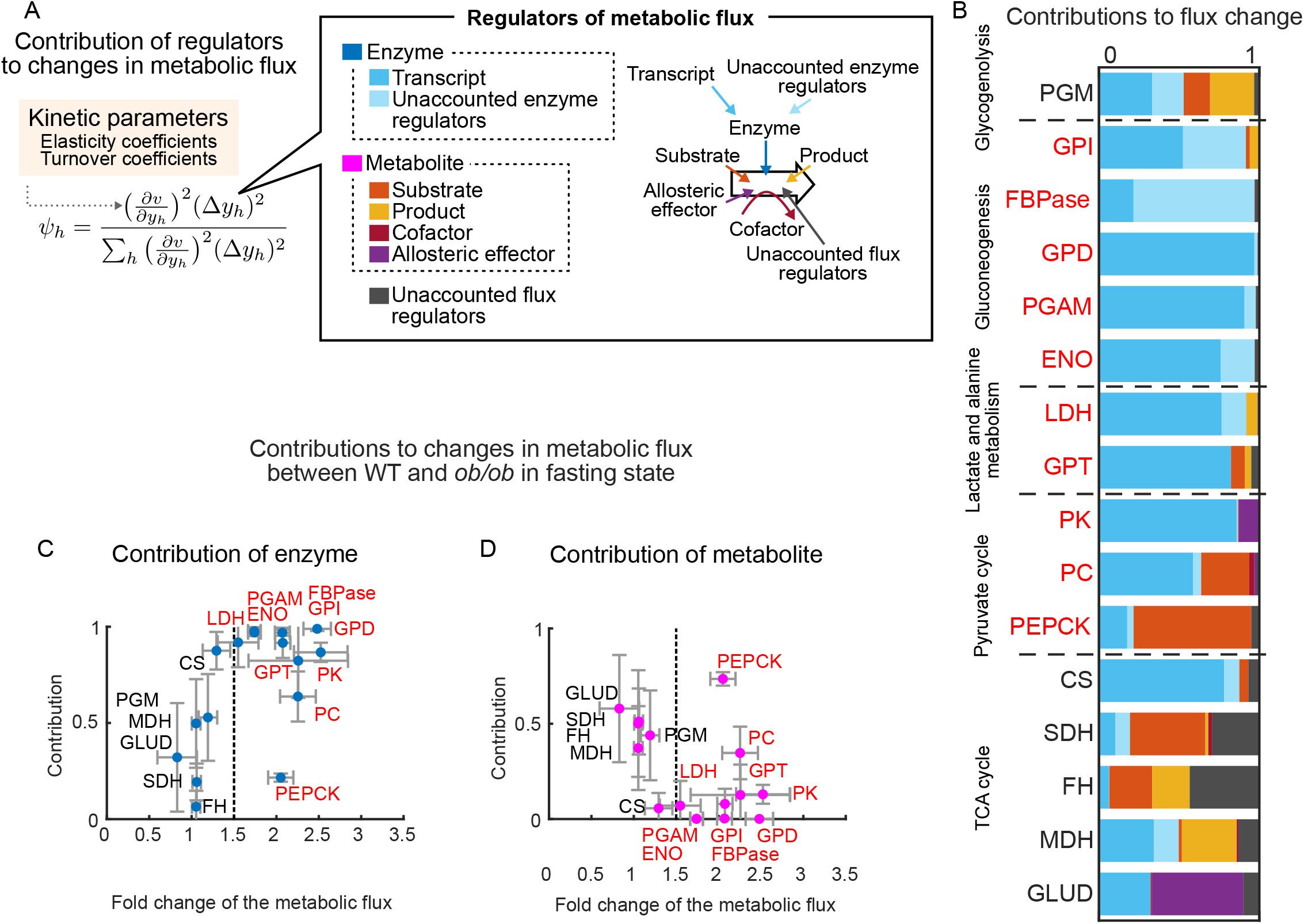
Contributions of regulators to changes in metabolic flux between fasting WT and *ob*/*ob*mice. (A) Schematic representation of contributions of regulators to changes in metabolic flux between conditions. The contribution was defined based on the sensitivity of the metabolic flux to the regulator, which is calculated using the kinetic parameters including elasticity coefficients (Figure S8; Table S5) and turnover coefficients (Table S5), and changes in the amounts of regulators between the conditions. See also Materials and Methods. (B) Contribution of regulators to changes in metabolic flux between WT and *ob*/*ob* mice in the fasting state. The reactions with the fold changes of the metabolic flux of *ob*/*ob* mice over that of WT mice in the fasting state larger than 1.5 are in red text. The stacked bars indicate the mean of the contributions independently calculated in all the Markov chain Monte Carlo samples in Figure S7. See also Table S6. (C, D) Scatter plots illustrating the relationships between the contributions of enzyme (C) or metabolite (D) to changes in metabolic flux and the fold changes of the metabolic flux of *ob*/*ob* mice over that of WT mice in the fasting state. For each reaction, the mean ± SD of the distribution of the contributions of enzyme or metabolite to changes in metabolic flux (x-axis) is plotted against the mean ± SD of the distribution of the fold changes of the metabolic flux of *ob*/*ob* mice over that of WT mice in the fasting state (y-axis). The vertical gray dotted line indicates the boundary where a fold change equals 1.5. The reactions with the fold changes of the metabolic flux of *ob*/*ob* mice over that of WT mice in the fasting state larger than 1.5 are in red text. See also Tables S5 and S6.

We defined contributions of regulators to changes in metabolic flux (Figure 5A). The regulators of metabolic flux were transcripts, unaccounted enzyme regulators, substrates, products, cofactors, allosteric effectors, and unaccounted flux regulators. Transcripts represent the mechanism by which changes in gene expression regulate enzyme abundance; the unaccounted ‘enzyme’ regulators represent other non-transcriptional mechanisms that influence the amount of enzyme such as protein degradation and stability. The unaccounted ‘flux’ regulators include such mechanisms as phosphorylation of enzymes and unknown allosteric effectors that were not accounted or measured in OMELET. The contribution of regulator ℎ to a change in metabolic flux through each reaction *ψ*_ℎ_ was calculated as

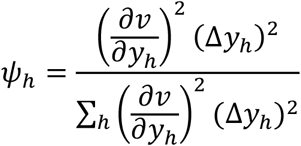

with a sensitivity of the metabolic flux to the regulator *v* (∂*v*/∂*y*_ℎ_) and a change in the amount of the regulator between conditions Δ*y*_ℎ_ . The contribution is calculated for changes in metabolic flux through each reaction between each pair of condition. The sum of the contributions of all regulators to a change in metabolic flux equals one, and a larger contribution indicates a stronger regulatory effect on metabolic flux. The contribution is a normalized value for each metabolic flux, thus is independent of the magnitude of the changes in metabolic flux between the conditions. The contribution was independently calculated in all the Markov chain Monte Carlo samples and represented as a distribution (Figure S7). We focused on the regulators with a mean contribution larger than 0.25; all regulatory contributions are available in Table S6.

We applied this analysis to evaluate the contribution of each type of regulator, including the unaccounted flux regulators, to difference in metabolic flux between WT and *ob*/*ob* mice in the fasting state (Figure 5B). We also examined the relationships between the contributions of enzymes or metabolites and the fold changes of the metabolic flux of *ob*/*ob* mice over that of WT mice in the fasting state (Figures 5C and 5D). For these analyses, the contribution of enzyme is the sum of its transcript and unaccounted enzyme regulators, and the contribution of metabolites is the sum of that of substrates, products, cofactors, and allosteric effectors. We found that, except for PEPCK, enzymes in the reactions with a fold change in metabolic flux greater than 1.5 in the *ob*/*ob* mice exhibited a greater contribution (Figure 5C) than did metabolites (Figure 5D). For PEPCK, the contribution of the enzyme was smaller and that of metabolites was larger. In particular, the substrate contributed the greatest effect (median: 0.73) on the increased metabolic flux through PEPCK (Figures 5B and S7; Table S6).

### Quantitative trans-omic networks for changes in metabolic flux between WT and *ob*/*ob* mice in the fasting state

To reveal a global landscape of alteration and dysregulation of metabolic flux in fasting *ob*/*ob* mice, we constructed a quantitative trans-omic network by integrating the experimental data (Figure 2), the changes in metabolic flux (Figure 4), and the contributions of the regulators (Figure 5). The resulting network consisted of four layers (Transcript, Enzyme, Metabolic Flux, and Metabolite) (Figures 6A-D). Nodes in Transcript, Enzyme, Metabolic Flux, and Metabolite layer represented the transcripts, enzymes, reactions, and metabolites. Lines connecting nodes in the Transcript layer to those in the Enzyme layer represented regulation of enzyme by transcript, and those by unaccounted enzyme regulators were not displayed. Lines connecting nodes in the Enzyme layer to reactions in the Metabolic Flux layer represented the contributions of the enzyme to changes in the metabolic flux between the conditions. Lines connecting nodes in the Metabolite layer to the reactions in the Metabolic Flux layer represented regulation of changes in metabolic flux by metabolites and were color-coded according to substrate, product, cofactor, or allosteric effector. Unaccounted flux regulators were not displayed. The size of nodes represents fold changes of the corresponding molecules or reactions in *ob*/*ob* mice over those of WT mice, and the width of the lines between the layers represents the contributions of regulators to changes in metabolic flux.

**Figure 6.**
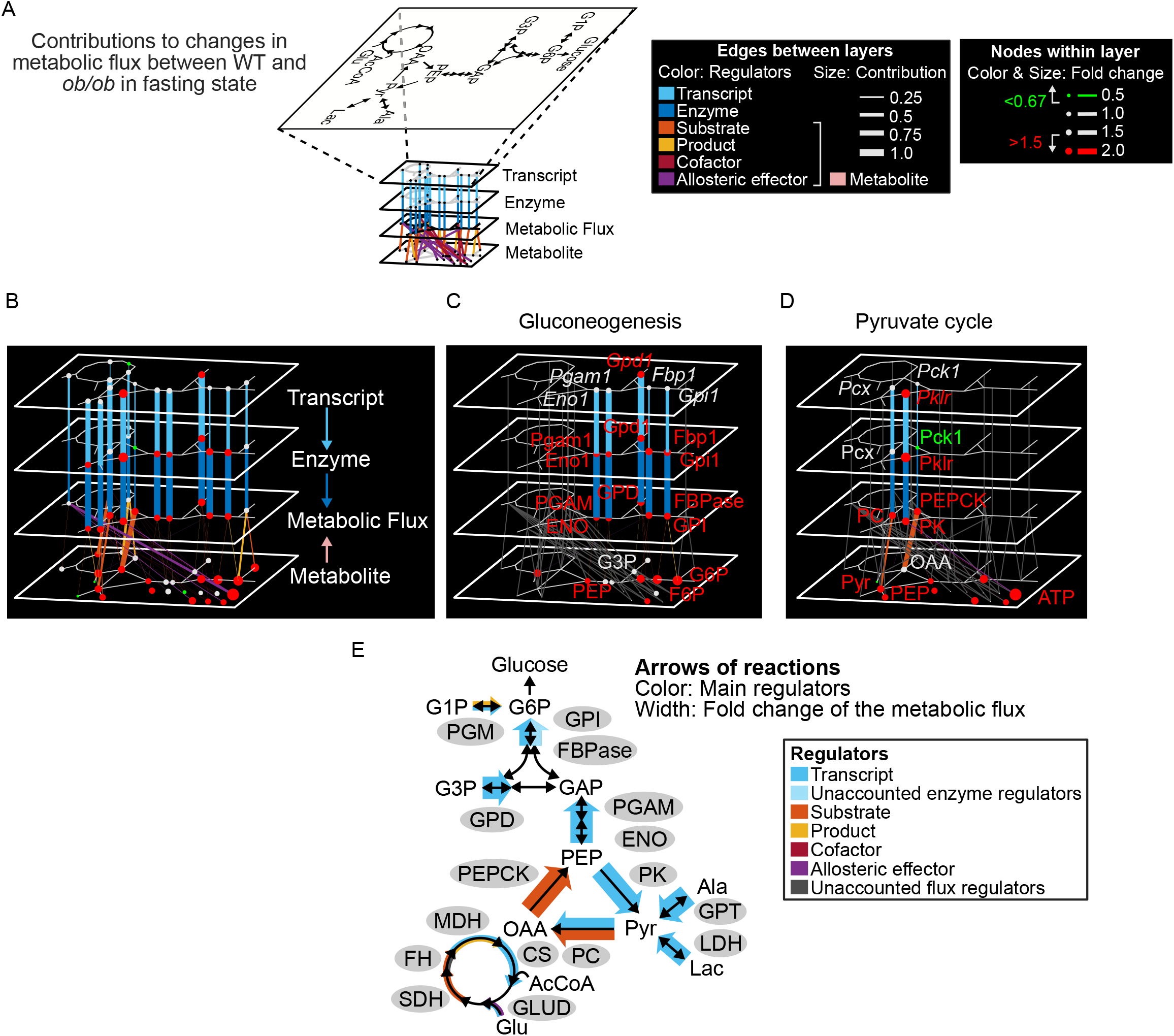
Quantitative trans-omic networks for changes in metabolic flux between WT and *ob*/*ob*mice in the fasting state. (A) Key to the quantitative trans-omic network for the difference between WT and *ob*/*ob* mice in the fasting state. See also Tables S2, S5, and S6. (B) The full quantitative trans-omic network. (C) The subnetwork of gluconeogenesis. (D) The subnetwork of pyruvate cycle. (E) A simplified metabolic pathway with metabolic fluxes and the contributions of main regulators. The color of the arrow in each reaction indicates the main regulators, which we defined as those with a mean contribution larger than 0.25. The size of the arrow in each reaction indicates fold changes of the metabolic flux in *ob*/*ob* mice over those in WT mice.

We extracted the subnetworks comprised of gluconeogenesis (Figure 6C) and of the pyruvate cycle (Figure 6D), which together represented the network with median of the fold changes of the metabolic flux of *ob*/*ob* mice over that of WT mice that were larger than 1.5 (Figure 5B, red text). In the subnetwork of gluconeogenesis (Figure 6C), many transcripts, enzymes, and metabolites also increased in *ob*/*ob* mice (2.2-fold increase in metabolites, 1.9-fold in enzymes, and 1.4-fold in transcripts on average within each layer) and size of nodes were qualitatively similar among Transcript, Enzyme, and Metabolite layers. By contrast, as for edges of contribution from one layer to another, the contributions of enzymes to metabolic flux were larger than those of metabolites. On average within the subnetwork, contribution of enzyme was 0.67 whereas contribution of metabolites was 0.017. In addition, contributions of transcripts to enzymes were similar among many enzymes including glycerol-3-phosphate dehydrogenase 1 (Gpd1), phosphoglycerate mutase 1 (Pgam1), and enolase 1 (Eno1) except for glucose-6-phosphate isomerase 1 (Gpi1) and fructose bisphosphatase 1 (Fbp1). These results indicate a hierarchical and quantitative regulation in gluconeogenesis and lactate and alanine metabolism, where 1.4-fold increase in transcripts contributed to 79% of 2.0-fold increase in metabolic fluxes while 2.2-fold increase in metabolites contributed to only 1.7% of the increase in metabolic fluxes (Figure 6E).

Pyruvate cycle consists of three reactions: PK reaction catalyzed by pyruvate kinase (Pklr) enzyme with the substrate PEP, PC reaction catalyzed by pyruvate carboxylase (Pcx) enzyme with the substrate pyruvate, and PEPCK reaction catalyzed by phosphoenolpyruvate carboxykinase 1 (Pck1) enzyme with the substrate oxaloacetate. Although fold changes in metabolic fluxes (2.5-fold for PK, 2.2-fold for PC, and 2.0-fold for PEPCK) and metabolites (1.5-fold for PEP, 1.9-fold for pyruvate, and 1.5-fold for oxaloacetate) in the subnetwork of pyruvate cycle were similar, changes in enzymes were different among reactions (Figure 6D): the enzyme Pklr increased, Pcx did not significantly changed, and Pck1 decreased in *ob*/*ob* mice. These differences in fold changes of molecules resulted in different contribution to metabolic flux among PK, PC, and PEPCK. Contribution of the enzyme Pklr to the PK flux (median: 0.87) was much larger than that of the substrate PEP (median: 0.0017) and Pklr enzyme mainly caused increase in PK flux in *ob*/*ob* mice. The medians of the contribution of Pcx enzyme and pyruvate to PC flux was 0.62 and 0.31, respectively, and both the enzyme and substrate contributed to increase in PC flux. Contribution of Pck1 enzyme to PEPCK flux (median: 0.21) was smaller than that of the substrate oxaloacetate (median: 0.74) and the substrate, rather than the enzyme, contributed increase in PEPCK flux. In all metabolic fluxes through PK, PC, and PEPCK in pyruvate cycle, contributions of *Pklr*, *Pcx*, and *Pck1* transcripts to Pklr, Pcx, and Pck1 enzymes (medians: 0.86, 0.59, and 0.18) were almost equal to those of Pklr, Pcx, and Pck1 enzymes to PK, PC, and PEPCK fluxes (medians: 0.87, 0.62, and 0.21), respectively, indicating that the contributions of the enzymes to changes in the metabolic flux were explained by those of the transcripts. These results suggested that the increased *Pklr* expression triggered the increased metabolic flux through the pyruvate cycle, which caused the accumulations of substrates, including pyruvate and oxaloacetate, and the large contributions of substrates to increases in metabolic fluxes through the downstream reactions of PC and PEPCK.

We also constructed a quantitative trans-omic network after oral glucose administration (Figure S9), which showed similar features with those in the fasting state. Thus, oral glucose administration had little effect on the differences in the steady-state metabolic flux between WT and *ob*/*ob* mice, and on the contributions of the regulators to the differences in the metabolic flux.

### Contributions of regulators to changes in metabolic flux induced by oral glucose administration within WT or *ob*/*ob* mice

We quantified the contributions of the regulators to changes in the metabolic flux by oral glucose administration in WT and *ob*/*ob* mice separately (Figures 7 and S7; Table S6). We examined the relationships between contributions of enzyme or metabolite to changes in the metabolic flux and fold changes of metabolic fluxes after oral glucose administration over those in the fasting state within each genotype (Figure 7A). None of the reactions showed a fold change of metabolic flux more than 1.5 nor less than 0.67, and no apparent relationship was found between the contributions of enzyme or metabolite and fold changes of the metabolic flux by oral glucose administration.

**Figure 7.**
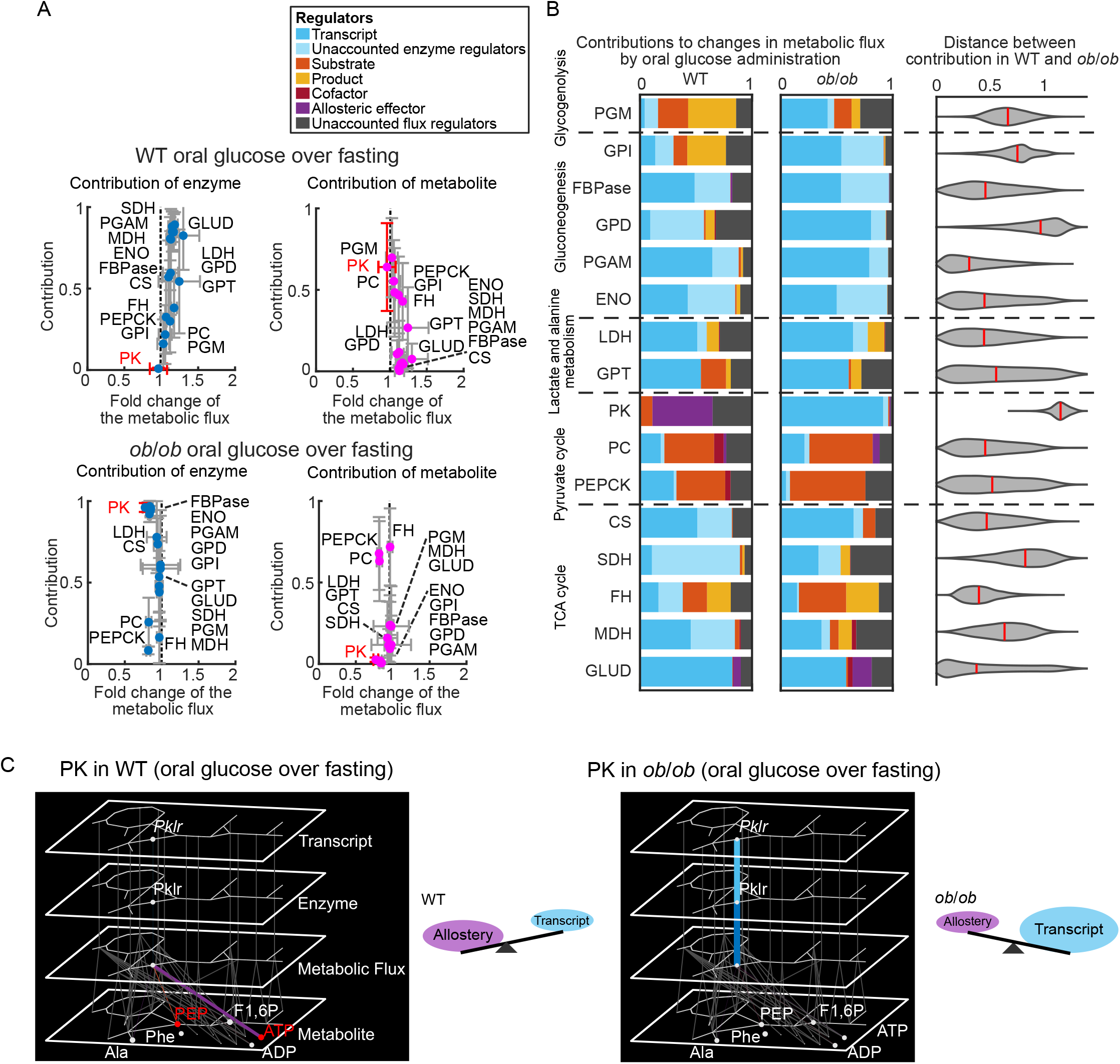
Contributions of regulators to changes in metabolic flux through PK by oral glucose administration. (A) Scatter plots illustrating the relationships between the contributions of enzymes and metabolites to changes in metabolic flux and the fold changes of the metabolic flux after oral glucose administration over that in the fasting state in WT mice (upper graphs) and *ob*/*ob* mice (lower graphs). For each reaction, the mean ± SD of the distribution of the contributions of enzyme or metabolite to changes in metabolic flux (x-axis) is plotted against the mean ± SD of the distribution of the fold changes of the metabolic flux of *ob*/*ob* mice over that of WT mice in the fasting state (y-axis). The vertical gray dotted line indicates the boundary where a fold change equals one. PK is highlighted in red. (B) Contribution of regulators to changes in metabolic flux between fasting and after oral glucose administration in WT mice (left) and *ob*/*ob* mice (middle). The stacked bars indicate the mean of the contributions independently calculated in all the Markov chain Monte Carlo samples in Figures S7. The violin plot in each reaction represents the distribution of the distance quantified as L2 norm between the contribution in WT and *ob*/*ob* mice independently calculated in all the Markov chain Monte Carlo samples. The vertical red line in each violin plot means the median of the distribution. See also Table S6. (C) Quantitative trans-omic networks for changes in metabolic flux through PK by oral glucose administration in WT mice and *ob*/*ob* mice. The networks had the same structures as those in Figure 6. Schematic representation of the main regulators to changes in metabolic flux between fasting and after oral glucose administration is displayed to the right of each network. We considered F1,6P as an allosteric activator, and ATP, alanine, and phenylalanine as allosteric inhibitors for PK in the Metabolite layer.

Among all the reactions, the largest difference in the contribution of the regulators by oral glucose administration between WT and *ob*/*ob* mice was found in PK (Figure 7B, right), a reaction with unchanged metabolic flux in WT mice and slightly decreased metabolic flux in *ob*/*ob* mice (Figure 4B). Allosteric effectors had the largest contribution to the change in metabolic flux through PK in WT mice (median: 0.55, Figures 7B and S7), whereas the *Pklr* transcript had the largest contribution in *ob*/*ob* mice (median: 0.95). These results suggested that the change in metabolic flux through PK by oral glucose administration was caused by different mechanisms between WT and *ob*/*ob* mice: changes in allosteric effectors in WT mice and changes in *Pklr* gene expression in *ob*/*ob* mice.

To explore the differences of the contributions of the regulators to the decreased metabolic flux through PK between WT and *ob*/*ob* mice, we constructed a quantitative trans-omic network for the change in metabolic flux through PK by oral glucose administration in WT mice (Figure 7C, left) and *ob*/*ob* mice (Figure 7C, middle). In WT mice, the substrate PEP and the allosteric inhibitor ATP in the Metabolite layer increased in response to oral glucose administration. The regulatory input from ATP in the Metabolite layer to PK in the Metabolic Flux layer (median: 0.50) was the largest among the regulatory inputs from the metabolites. In *ob*/*ob* mice, no metabolites in the Metabolite layer increased nor decreased following oral glucose administration. The regulatory input from Pklr in the Enzyme layer to PK in the Metabolic Flux layer (median: 0.95) was larger than regulatory inputs from metabolites, including PEP, ADP, fructose 1,6-bisphosphate (F1,6P), ATP, alanine, and phenylalanine, each of which had a regulatory input less than 0.10. The regulatory input from *Pklr* in the Transcript layer to Pklr in the Enzyme layer (median: 0.93) was almost equal to that from Pklr in the Enzyme layer to PK in the Metabolic Flux layer (median: 0.95), indicating that the contribution of enzyme was explained by that of transcript. These results suggested that the change in metabolic flux through PK was caused by increased ATP as an allosteric inhibitor in WT mice and by slightly decreased *Pklr* transcript in *ob*/*ob* mice. Given that the glucose-induced changes in metabolic flux through PK were not large (Figure 4B), we interpreted these findings to indicate that WT and *ob*/*ob* mice used different regulatory mechanisms, allosteric regulation and transcripts, respectively, to maintain the metabolic flux through PK rather than to change metabolic flux.

## DISCUSSION

In this study, we developed a method OMELET to investigate alterations and dysregulation of metabolic flux in liver that are associated with obesity. Using OMELET, we inferred the metabolic fluxes in glucose metabolism in livers of WT and *ob*/*ob* mice in the fasting state and after oral glucose administration to identify changes in metabolic flux between the conditions. The metabolic flux through reactions in gluconeogenesis and the pyruvate cycle increased in *ob*/*ob* mice compared to WT mice in the fasting state. The increased metabolic fluxes through reactions in gluconeogenesis were mainly caused by increased transcripts. In the pyruvate cycle, increases in transcripts mediated the increased metabolic flux through PK and increases in substrates the increase through PEPCK. In response to oral glucose administration, differences in the metabolic fluxes within mice of the same genotype were small compared to those between WT and *ob*/*ob* mice. Oral glucose administration did not change metabolic flux through PK in either WT or *ob*/*ob* mice, but the metabolic flux was regulated by increased ATP in WT mice and by decreased *Pklr* transcript in *ob*/*ob* mice. Thus, WT and *ob*/*ob* mice used different regulatory mechanisms, allosteric regulation and transcripts, respectively, to maintain the metabolic flux through PK rather than to change metabolic flux.

Although isotopic labeling is a powerful technique to measure metabolic flux, the introduction of isotopic tracers into living animals may perturb the activity of the metabolic pathway of interest. OMELET does not require isotopic labeling data to infer metabolic flux, thus avoiding these potential perturbations caused by the addition of isotopic tracers. However, the accuracy of inference of metabolic flux without isotopic labeling data needs to be validated. We validated the performance of OMELET by applying it to the simulated datasets from a kinetic model of the yeast glycolysis (Messiha et al., 2014; Smallbone et al., 2013) (Figure S2). We also applied OMELET to the data in fasting mouse liver and found that the inferred metabolic fluxes in WT mice were consistent with those in the previous studies (Figures S5A and S5B). The fold changes of the metabolic fluxes through most reactions in the glucose metabolism of *ob*/*ob* mice over that of WT mice inferred by OMELET were consistent with those in the previous studies, except for that through glycogenolysis (Figures S5C and S5D). These results suggested that the experimental data of the amounts of enzymes and metabolites contain sufficient information on metabolic fluxes as latent parameters, which can be inferred by OMELET.

We used the simultaneously obtained experimental dataset of the amounts of metabolites, enzymes, and transcripts from the same samples for OMELET. Using simultaneously obtained multi-omic data, a Bayesian method has the potential to analyze metabolic flux (Heinonen et al., 2019) and its regulation (Hackett et al., 2016; John et al., 2019; Saa and Nielsen, 2016). Such datasets enabled us to apply a Bayesian method rather than analyzing the population mean. A Bayesian method can incorporate uncertainties inherent in the experimental data, such as measurement noise and population heterogeneity. Based on a Bayesian method, we assumed the experimental data resulted from a generative model that described the underlying processes given latent parameters, and evaluated the probability that the model yields the data by likelihood. In OMELET, the enzymes were derived from a generative model based on metabolic flux, and the transcripts from a generative model based on protein turnover (Figure 3). Using these two generative models in OMELET, we evaluated the likelihood of the enzymes and transcripts from each mouse to infer unknown parameters including metabolic fluxes, elasticity coefficients and protein turnover coefficients.

There are several limitations to this study. The 95% credible intervals of the metabolic fluxes in the TCA cycle were large compared to those through other reactions in glucose metabolism (Figures 4A and S3), indicating that the metabolic fluxes in the TCA cycle were not reliably determined. The inaccuracy of the inferred metabolic fluxes in the TCA cycle may be due to small changes in the amounts of enzymes and metabolites in the TCA cycle between the conditions. We did not consider the compartmentation of reactions into cytoplasm and mitochondria, and inferred metabolic fluxes averaged in a whole cell, which may result in the inaccuracy of the inferred metabolic fluxes. In addition, OMELET is based on a steady-state assumption and cannot infer dynamic changes in metabolic flux under non-steady-state conditions.

Using OMELET, we found altered and dysregulated metabolic flux associated with obesity. We found that the large increase in metabolic flux through reactions in gluconeogenesis in *ob*/*ob* mice compared to WT mice in the fasting state was mainly caused by increased gene expression of the enzymes (Figure 6). There are several transcription factors involved in controlling the expression of genes encoding enzymes involved in gluconeogenesis. For example, cAMP response element-binding protein (CREB) activates transcription of *G6pc* and *Fbp1* (Hanson and Reshef, 1997; Herzig et al., 2001), as well as *Gpi1* and *Pgam1* (Everett et al., 2013). Liver-specific knockdown of CREB reduced fasting plasma glucose concentrations in *ob*/*ob* mice through downregulation of *G6pc* and *Fbp1* (Erion et al., 2009). In addition to these key transcription factors identified by individual experiments, high-throughput measurements and multi-omic analyses have revealed many more transcription factors involved in metabolic alteration associated with obesity (Egami et al., 2021; Kokaji et al., 2020; Soltis et al., 2017). Although transcription factors contribute changes in metabolic flux in glucose metabolism associated with obesity, metabolic flux is also regulated by metabolites that include substrates, products, cofactors, and allosteric effectors. In this study, we found that transcripts, rather than metabolites, mainly contributed to the differences in the metabolic flux between WT and *ob*/*ob* mice (Figure 4). Our results suggested that transcription factors would trigger increased gluconeogenic flux associated with obesity by promoting the expression of the genes encoding the relevant metabolic enzymes.

In the pyruvate cycle, increased oxaloacetate (a substrate), rather than Pck1, contributed to the increased metabolic flux through PEPCK in fasting *ob*/*ob* mice (Figure 6D). Several metabolic flux analyses showed that the metabolic flux through PEPCK increased associated with obesity (Patterson et al., 2016; Satapati et al., 2012; Sunny et al., 2011), which was consistent with our data. Given that PEPCK is an irreversible reaction and there are no known allosteric effectors, possible regulators of the metabolic flux through PEPCK include Pck1 amounts, the substrate oxaloacetate, and the cofactor GTP. However, it has been unclear which regulator mainly contributed the increased metabolic flux through PEPCK associated with obesity. Among the possible regulators, Pck1 amounts decreased associated with obesity in the fasting state (Samuel et al., 2009; Satapati et al., 2012; Sunny et al., 2011), which was observed in our proteomic data (Figure 2). These studies suggest that the increased metabolic flux is not consistent with the decreased Pck1 amount. Here, we found that increased oxaloacetate was responsible for the increased metabolic flux through PEPCK (Figure 6D), which provides a mechanistic explanation for the increased metabolic flux through PEPCK that is associated with obesity.

The only difference in the contributions of regulators to the changes following oral glucose administration between WT and *ob*/*ob* mice was in metabolic flux through PK (Figure 7). In WT mice, the allosteric effector ATP was the largest contributor to the slightly decreased PK flux. In *ob*/*ob* mice, a reduction in *Pklr* transcript was the largest regulatory contributor. A reason why allosteric regulation was not the main regulator in *ob*/*ob* mice may be because amounts of allosteric effectors, such as ATP, were high even in the fasting state and did not increase following oral glucose administration (Figure 2).

In calculating the contributions of regulators to changes in metabolic flux between the conditions, we considered unaccounted flux regulators as one of the regulators of metabolic flux and unaccounted enzyme regulators as one of the regulators of enzymes. The contributions of unaccounted flux regulators to changes in metabolic flux between fasting WT and *ob*/*ob* mice were smaller than those of other regulators in all the reactions in gluconeogenesis and the pyruvate cycle, indicating that changes in metabolic fluxes through these reactions can be explained by known regulators that we considered (Figures 5B and S7). In contrast, the contributions of unaccounted enzyme regulators to changes in enzyme in reactions through GPI and FBPase were relatively larger than those through other reactions, indicating that the increase in enzyme in *ob*/*ob* mice cannot be explained by the transcripts. Although we used a simple linear relationship between the amount of an enzyme and a transcript in OMELET, an increase in enzyme that is not accompanied by the changes in transcripts should be explained by other regulatory mechanisms, such as changes in protein stability. Furthermore, the contributions of unaccounted flux regulators to changes in metabolic flux between fasting and after oral glucose administration were larger in many reactions than those between fasting WT and *ob*/*ob* mice (Figures 7B and S7). To explain the contributions of unaccounted flux regulators, we need to consider posttranslational modifications, such as phosphorylation, of enzymes. Such data can be incorporated by including phosphoproteomic data.

In conclusion, we developed OMELET, which uses the simultaneously obtained multi-omic data to infer metabolic fluxes in the glucose metabolism in multiple conditions and to identify changes in metabolic flux between the conditions. Furthermore, we quantified the contributions of the regulators to the changes in metabolic flux between the conditions. OMELET is designed to infer metabolic flux without using isotopic labeling data and to simultaneously infer changes in metabolic flux and the contributions of regulators. The quantitative trans-omic network provided insights into the obesity-associated changes in the glucose metabolism in liver and revealed comprehensive molecular mechanisms for understanding the pathology of alteration and dysregulation of metabolic flux associated with obesity.

## ACKNOWLEDGEMENTS

We are grateful to Maki Ohishi, Ayano Ueno, Keiko Endo, Sanae Ashitani, Keiko Kato, and Kaori Saitoh (Keio University) for their technical assistance with metabolomic analysis using CE-MS. We thank for Shinsuke Uda of Kyushu University and our laboratory members for critically reading this manuscript and for their technical assistance with the experiments. The computational analysis of this work was performed in part with support of the supercomputer system of National Institute of Genetics (NIG), Research Organization of Information and Systems (ROIS). This manuscript was edited by Nancy R. Gough (BioSerendipity, LLC).

This work was supported by the Japan Society for the Promotion of Science (JSPS) KAKENHI Grant Number JP17H06300, JP17H06299, JP18H03979, JP21H04759 and by The Uehara Memorial Foundation. S.U. receives funding from JSPS KAKENHI Grant Number JP19J22134. S.O. receives funding from a Grant-in-Aid for Early-Career Scientists (JP17K14864 and JP21K14467). A. Hatano is supported by Grant-in Aid from the Tokyo Biochemical Research Foundation. T.K. receives funding from JSPS KAKENHI Grant Number JP21K16349. H.K. was supported by JSPS KAKENHI JP20H03237. Y.S. was supported by the JSPS KAKENHI Grant Number JP17H06306. K.I.N. was supported by the JSPS KAKENHI Grant Number JP17H06301, JP18H05215. A. Hirayama was supported by the JSPS KAKENHI Grant Number JP18H04804. T.S. receives funding from the AMEDCREST from the Japan Agency for Medical Research and Development (AMED) under Grant Number JP18gm0710003.

## AUTHOR CONTRIBUTIONS

S.U., S.O., and S.K. conceived the project; A. Hatano, T.K., Y.I., H.K., M.M., and K.I.N. designed and performed the animal experiments; A. Hirayama and T.S. performed metabolomic analysis using CE-MS; Y.S. performed transcriptomic analysis using RNA-seq; M.M. and K.I.N. performed proteomic analysis using LC-MS/MS; S.U., S.O., K.T., A. Hatano, T.K., Y.I., and K.H. analyzed the data; S.U., S.O., and K.T. performed mathematical modeling analyses. S.U., S.O., and S.K. wrote the manuscript. All authors read and approved the final manuscript.

## DECLARATION OF INTERESTS

The authors declare no competing interests.

## MATERIALS AND METHODS

### RESOURCE AVAILABILITY

#### Lead Contact

Further information and requests for resources and reagents should be directed to and will be fulfilled by the Lead Contact, Shinya Kuroda.

#### Materials Availability

This study did not generate new unique reagents.

#### Data and Code Availability

The datasets generated during this study are in the published article. The MATLAB and R code for OMELET is available at GitHub (https://github.com/usa0ri/OMELET). An image for Docker container that include RStan and R software to perform OMELET is available at DockerHub Registry (https://hub.docker.com/repository/docker/saori/rstan). The accession number for the data of proteome analysis reported in this paper is the ProteomeXchange Consortium (http://proteomecentral.proteomexchange.org) via the JPOST partner repository: JPST000147, JPST000148, and JPST001222. Sequence data used in this study have been deposited in the DNA Data Bank of Japan Sequence Read Archive (DRA) (https://www.ddbj.nig.ac.jp/) under the accession no. DRA008416 and no. DRA012292.

## EXPERIMENTAL MODEL AND SUBJECT DETAILS

### Animals

We used the mouse liver samples obtained simultaneously with those previously published (Egami et al., 2021; Kokaji et al., 2020). Briefly, 10-week-old male C57BL/6J wild-type and *ob*/*ob* mice (Japan SLC, Inc., Shizuoka, Japan) were overnight (16 hours) fasted or administrated 2 g/kg body weight of glucose orally after overnight fasting. The mice before or four-hour after glucose loading were sacrificed by cervical dislocation and the whole or left lobe of the liver was dissected and immediately frozen in liquid nitrogen. The frozen liver was pulverized with dry ice to a fine powder with a blender and separated into tubes for transcriptomic, proteomic, and metabolomic measurements. Note that all the omics data was simultaneously measured from the same individual mice. All the mouse experiments were performed according to protocols approved by the animal ethics committee of The University of Tokyo.

We had mice under four conditions: WT mice in the fasting state (n=11), *ob*/*ob* mice in the fasting state (n=12), WT mice after oral glucose administration (n=12), and *ob*/*ob* mice after oral glucose administration (n=12). The metabolomic data of five samples in each condition were reported in our previous studies (Egami et al., 2021; Kokaji et al., 2020). We newly obtained the metabolomic data from all the samples in this study. The transcriptomic data from all the samples in the fasting state and five after oral glucose administration were reported in our previous studies. We newly obtained the transcriptomic data from seven samples after oral glucose administration in this study.

## METHOD DETAILS

### Metabolic network for glucose metabolism in mice

A metabolic network for glucose metabolism in mice was constructed to infer metabolic fluxes and quantify the contributions of regulators to changes in metabolic flux between conditions. The network consists of 27 metabolites and 22 reactions (Table S1) in gluconeogenesis, glycogenolysis, lactate and alanine metabolism, and the TCA cycle (Figure 1).

The liver produces glucose through the gluconeogenesis and consumes glucose through glycolysis. The gluconeogenesis in liver includes reactions catalyzed by glucose-6-phosphate (G6PC) and fructose-bisphosphatase 1 (FBPase), while the glycolysis includes reactions by glucokinase (GK) and phosphofructokinase (liver type) (PFKL).

Assuming that the metabolic flux through GK and PFKL were negligible in livers of WT and *ob*/*ob* mice in all the conditions, we included G6PC and FBPase in the metabolic network but not included GK and PFKL. This is supported by several studies showing that the glucose production was more dominant than the glucose consumption in livers of overnight fasting WT mice (Burgess et al., 2005; Hasenour et al., 2015, 2020; Satapati et al., 2012; Wang et al., 2020) and *ob*/*ob* mice (Turner et al., 2005). In addition, the glucose production from glycogenolysis and gluconeogenesis occurred 150 min after oral glucose administration in rat (Jin et al., 2003). Although we did not find any studies to support the glucose production in *ob*/*ob* mice four hours after oral glucose administration, *ob*/*ob* mice after oral glucose administration in this study showed similar temporal changes in blood glucose and insulin (Figures S1A and B) to WT mice. Therefore, we included G6PC and FBPase in the metabolic network but not included GK and PFKL reactions. This also reduces the complexity of the metabolic model and computational cost for the metabolic flux estimation.

Pyruvate dehydrogenase (PDH), converting pyruvate to acetyl-CoA, was not included in the metabolic network because several studies showed that the metabolic flux through PDH was small (~5%) relative to those through the TCA cycle in WT mice in the fasting state (Perry et al., 2016). Malic enzyme (ME), converting malate to pyruvate, was not considered because ME inhibitor did not affect the metabolic flux producing pyruvate in fasting rodent models (Hasenour et al., 2020; Perry et al., 2016; Petersen et al., 1995), suggesting the small contribution of ME to the metabolic flux.

Cytoplasmic and mitochondrial compartments were not considered for simplification and averaged metabolic fluxes as a single compartment were inferred in this study. Malate dehydrogenase (Mdh) has cytoplasmic (Mdh1) and mitochondrial (Mdh2) isoforms, but we only considered Mdh2 as a part of reactions in the TCA cycle for simplification.

### Algorithm of omics-based metabolic flux estimation without labeling for extended trans-omic analysis (OMELET)

We first review a framework of Bayesian inference. Statistical model consists of a likelihood, representing a probability of the observed data ***z*** at a given parameter values ***θ***, and a prior distribution for ***θ***, denoting a probability distribution of each parameter reflecting the feasible assumptions and prior knowledges. Bayes’ theorem calculates the renormalized product of the likelihood *p*(***z***|***θ***) and the prior distribution *p*(***θ***),

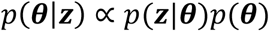

to produce the posterior parameter distribution *p*(***θ***|***z***), a probability of the parameters taking the values given the observed data. The posterior distribution is obtained by updating the parameter values following the prior distributions toward better fittings to the observed data evaluated in likelihood. The definition of the prior distribution is critical especially when the sample size is small whereas sufficient samples make its effects on posterior parameter distribution decreases to get closer to maximal likelihood estimation. One may assume additional parameters for prior distributions, which can be achieved by defining hyperparameters ***η***,

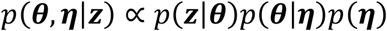

to introduce a hierarchical structure in prior distributions. Furthermore, Bayesian regression uses the additional observed data ***w*** as explanatory variables to calculate posterior distributions defined as

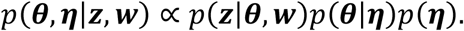

In Omics-based Metabolic flux Estimation without Labeling for Extended Trans-omic analysis (OMELET), observed data ***z*** is the experimental data of the amounts of enzymes and transcripts from the same samples obtained from multiple conditions. The observed data ***w*** as explanatory variables is the experimental data of the amounts of metabolites. Parameters ***θ*** include metabolic fluxes in each condition, elasticity coefficients in linlog kinetics and protein turnover coefficients. The prior *p*(***θ***) is defined for all the parameters, and the hyperprior *p*(***η***) is defined only for metabolic fluxes, which has hyperparameters ***η*** to obtain a steady-state metabolic flux distribution based on the reactions of the metabolic network of interest. The likelihood *p*(***z***|***θ***), which describes the relationship between the experimental data and the parameters, is the product of the probability of the measured amounts of enzymes given the parameters and the probability of the measured amounts of transcripts given the parameters.

We start from defining the prior and hyperprior for metabolic fluxes as multivariable normal distribution in each condition *l* = 1,2, … , *g* . The metabolite concentration ***x***_*l*_ and metabolic fluxes ***v***_*l*_ describes a system of mass balances around each metabolite in the form

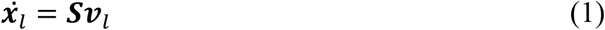

for *r*′ reactions and *m* metabolites, where ***S*** denotes the stoichiometric matrix that links metabolites to their reactions via stoichiometry. Note that this stoichiometric matrix is based on the open-formed metabolic network that is transformed by removing rows for the metabolites that participated in transporting reaction across the system boundary, resulting in *m* < *r*′. The vector of the time derivative for metabolites 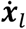 around steady state is assumed to follow *m*-dimensional multivariate normal distribution

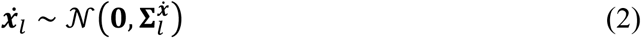

with diagonal covariance matrix 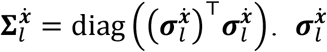 represents the extent of relaxation from the steady state and is defined later based on the influx and efflux around each metabolite. The number of variables that need to be specified to calculate a steady-state fluxes in equation (1) is *f* = *r*′ − rank(***S***). Let us denote the vectors of independent and dependent flux variables ***u***_*l*_ and 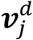 of length *f* and rank(***S***), respectively. Since dependent flux variables can be directly computed as linear combination of independent flux variables, we have only to estimate independent flux as hyperparameter to obtain the full metabolic fluxes. Here the vector of independent flux was assumed to follow the multi-dimensional normal distribution

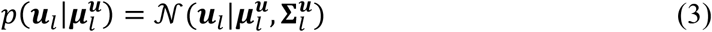

with mean 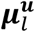 and diagonal covariance matrix 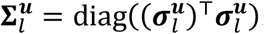. The deviations of independent fluxes were determined as 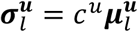 with a fixed coefficient of variance *c*^*u*^. The relation between independent and dependent flux variables can be obtained via decomposition of the full flux vector into the vectors of independent and dependent flux, and the equation (1) can be expressed as

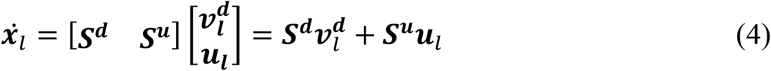

where *m* × rank(***S***) matrix ***S***^*d*^ and *m* × *f* matrix ***S***^*u*^ contain columns corresponding to dependent and independent flux variables, respectively. When ***S***^*d*^ is regular with *m* = rank(***S***) and det(***S***^*d*^) ≠ 0, the full flux vector is directly computed as

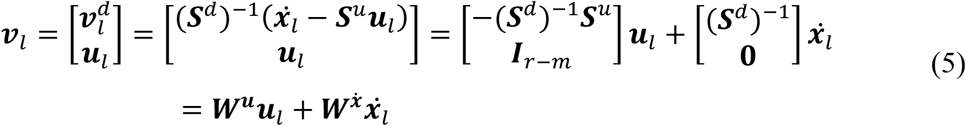

with transformation matrices ***W***^***u***^ and 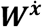 defined based on the inverse of ***S***^*d*^. Here we just considered the metabolic pathways in which the stoichiometric matrix ***S***^*d*^ was regular, and in other words the following situation was not considered because the full metabolic fluxes could not be computed from independent flux variables: *r*′ < *m* or linearly dependent rows in ***S*** resulting in rank(***S***) < *m*. We avoided these conditions by appropriate definition of the target metabolic pathway and stoichiometric matrix.

Now we obtain the prior distribution of the full metabolic fluxes from equation (5). Since linear combination of normal random variables is also normal random variables, the vector of full metabolic flux follows the *r*′-dimensional normal distribution

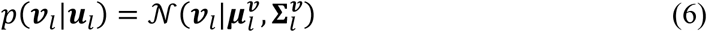

with mean 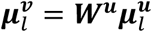 and covariance matrix 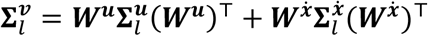. The diagonal covariance matrix 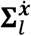 represents the variances around prior metabolite changes. In strict steady state, the prior for metabolite change becomes Dirac’s delta function at zero by increasing the variances, we can relax the steady-state assumption on individual metabolites and encode allowance for accumulations or depletions of them. The squared diagonal element 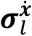 was obtained from the mean of the prior distribution of metabolic flux as

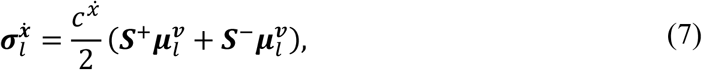

with a fixed coefficient of variance 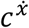 by defining production and consumption stoichiometric matrices as,

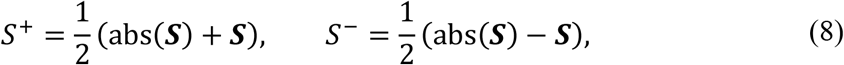

where abs(***S***) is the matrix of absolute values of the corresponding entries of ***S***. Note that the entries of the matrix ***S***^+^ corresponds to the number of molecules of metabolite produced by the reaction. Conversely, each entry of the matrix ***S***^−^ give the number of molecules of metabolite consumed by the reaction.

Next, we define the likelihood of the measured amounts of enzymes given parameters and prior for elasticity coefficients based on linlog kinetics. Since the experimental data of the amounts of metabolites and enzymes is usually not available for all reactions in the metabolic network, the likelihood was calculated for a subset of the reactions. We consider the metabolic flux through the subset of the reactions *R* ⊆ *R*′ = {1,2, … , *r*′}, where *R* consists of *r* reactions. For each sample *k* = 1,2, … , *n*_*j*_ under condition *l* (*l* = 1,2, … , *g*), the amounts of metabolites ***x***_*kl*_ (a *m* × 1 vector) and the amounts of enzymes ***e***_*kl*_ (a *r* × 1 vector) are obtained after normalizing by the average amounts across all the conditions. In the linlog kinetics framework, the metabolic flux through reaction 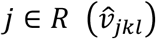 is expressed as

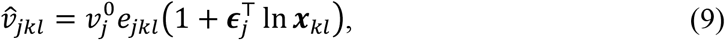

where 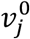 is the metabolic flux in the reference state, and ***ϵ***_*j*_ is the *m* × 1 vector of elasticity coefficients. 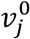 is defined as the mean of the prior metabolic flux values across conditions as 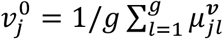 where 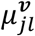 is a prior mean of the metabolic flux *v*_*jl*_. *ϵ*_*ji*_ describes the effect of changes of the amounts of metabolites *x*_*i*_ on the metabolic flux *v*_*j*_, and is positive if metabolite *i* is a substrate or an allosteric activator for reaction *j*, while negative if the metabolite is a product or an allosteric inhibitor. If metabolite *i* does not directly participate in reaction *j*, the value of *ϵ*_*ji*_ equals to zero. According to equation (9) the amount of the enzyme is calculated using the inferred metabolic flux *v*_*jl*_ . Here, the amount of the enzyme in each sample *e*_*jkl*_ is assumed to follow the normal distribution around the estimated value 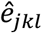, and we obtain the likelihood of the measured amount of the enzyme given parameters as

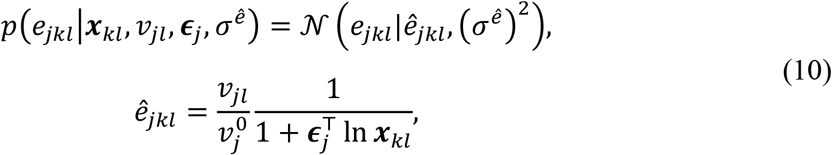

where *v*_*jl*_ is the inferred metabolic flux of reaction *j*. For simplicity, the variance of the error term 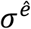 is set to the same values in all the reactions, samples, and conditions. We placed half-Cauchy priors with scale 0.5 on 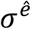, set as a weakly informative prior distribution given that 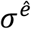 was expected to be less than one.

Elasticity coefficients are likely not to significantly deviate from the range between −1 and 1 theoretically (Kacser and Burns, 1995), and are likely to be positive for substrates and allosteric activators whereas negative for products and allosteric inhibitors. This property generates prior distributions for elasticity coefficients as

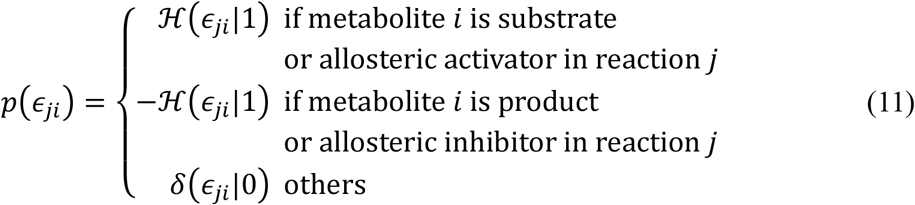

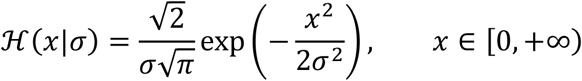

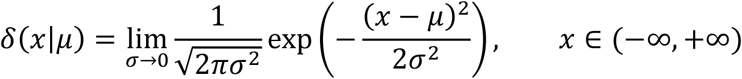

where 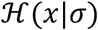 indicates half-normal distribution with variance *σ* , and *δ*(*x*|*μ*) indicates Dirac’s delta function equal to zero everywhere except for *μ*.

The amounts of enzymes are explained in the context of not only metabolic flux but also of protein turnover. Here we define the likelihood of the measured amounts of transcripts given parameters and priors for protein turnover coefficient *β*_*ij*_ . For each sample *k* = 1,2, … , *n*_*j*_ under condition *l* (*l* = 1,2, … , *g*) , the estimated amounts of enzymes 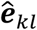 and the corresponding amounts of transcripts 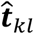 represents the enzyme change rate 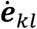 as

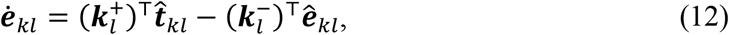

where 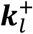 and 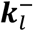 are *r* × 1 vectors of kinetic parameters for protein synthesis and degradation, respectively. Assuming the amounts of enzymes as stable within the observed time intervals, the amounts of enzymes and transcripts have a linear relationship and we obtain the likelihood of the measured amount of the transcript given parameters as,

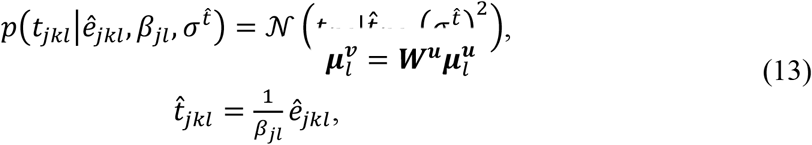

using turnover coefficient 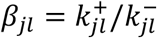 and the estimated amount of enzyme 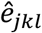 in each sample defined in equation (10). The parameter to determine the variance of the error term 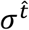 is simplified as the common values in all the reactions, samples, and conditions. We placed half-Cauchy priors with scale 0.5 on 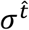. Given that there is a high correlation between copy numbers of RNA and protein especially in the glucose metabolism (Matsumoto et al., 2017), the turnover coefficient *β*_*jl*_ is expected to be close to one when the amounts of enzymes and transcripts are normalized to their averages. Therefore, the prior distribution for *β*_*jl*_ can be described by

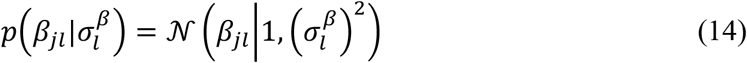

with error term defined by 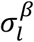 in each condition. We placed half-Cauchy priors with scale 0.5 on 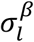.

Combining the likelihood with the priors based on linlog kinetics (equations 6, 10, 11, and Table S7) and protein turnover (equations 13, 14, and Table S7), the joint posterior distribution is given by

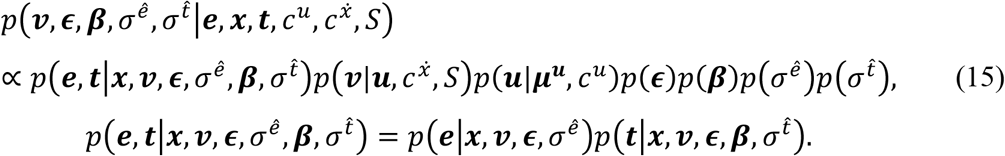

where 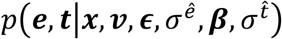 is the likelihood of the measured amounts of enzymes based on linlog kinetics combined with the likelihood of the measured amounts of transcripts based on protein turnover, 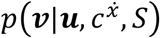 is the prior distribution for metabolic flux ***v***,*p*(***u***|***μ***^***u***^, *c*^*u*^) is the hyperprior distribution for independent flux ***u***,*p*(***ϵ***) is the prior distribution for elasticity coefficients ***ϵ***, *p*(***β***) is the prior distribution for protein turnover coefficients ***β***, 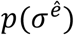 is the prior distribution for parameter of error term 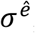, and 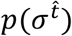 is the prior distribution for parameter of error term 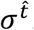.

### Application of OMELET to mouse data

We applied OMELET to the experimental data from mice in four conditions: WT in the fasting state, WT after oral glucose administration, *ob*/*ob* in the fasting state, and *ob*/*ob* after oral glucose administration. The independent fluxes were constrained so that the flux through G6PC in WT mice in the fasting state was fixed at one. A metabolic flux through each reaction was inferred simultaneously in all the conditions and as inferred as the relative value to that through G6PC in WT mice in the fasting state. The amounts of pyruvate and oxaloacetate were not available in our measurements because of their chemical instability and low concentrations. For such metabolite species, the relative amounts normalized to the mean across the conditions were inferred as parameters. The amounts of enzymes of glutamic pyruvic transaminase (Gpt) and glutamate dehydrogenase 1 (Glud1) were not measured and only the likelihood of the measured amounts of transcripts was evaluated. Several enzymes function as complex including succinate dehydrogenase, which is also known as respiratory complex II. Two subunits of succinate dehydrogenase, subunit A (Sdha) and subunit B (Sdhb), were available both in the amounts of enzymes and transcripts. The amounts of Sdha and Sdhb were independently normalized to the mean values across the conditions, and then the product was introduced as ***e*** for the reaction through succinate dehydrogenase (SDH) in equation (10). Other reactions were catalyzed by a single enzyme or by complex with only one subunit measured and a single enzyme data was used. The parameters for coefficient of variances *c*^*u*^ and 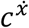 were fixed at 0.1 and 0.01, respectively.

We performed posterior predictive checks to evaluate the fitting of the model to the measured data of the amounts of enzymes and transcripts from mice. Posterior predictive distributions of enzymes can be simulated by sampling parameters from the posterior and using them to generate replicate data sets based on equation (10). Posterior predictive distributions of transcripts can be simulated in the same way based on equation (13). We compared the posterior predictive distributions with the measured data and confirmed the good fits to the experimental data of the amounts of enzymes and transcripts (Figure S4).

### Parameter estimation

Based on the specified prior distribution and likelihood, the posterior distributions of parameters were numerically estimated by Markov Chain Monte Carlo (MCMC) sampling. The algorithm was a No-U-turn sampler (NUTS), a variant of Hamiltonian Monte Carlo (HMC), constructing an iterative process that eventually converges to the true posterior distribution (Hoffman and Gelman, 2014). For application to the data from mice, we ran four chains of 20,000 iterations with 10,000 burnings with thinning of 2, resulting in 20,000 samples in total. Convergence of Markov Chains was evaluated by R-hat diagnostic, which compares the between- and within-chain estimated for model parameters. All our runs satisfied R-hat less than 1.05, indicating that chains were mixed well. All the parameter estimation was performed using RStan library (version 2.19.2) in R software (3.6.1) within a Docker container (Merkel, 2014).

### Simulation using kinetic model of yeast glycolysis

The model of yeast glycolysis was downloaded from the public model repository BioModels Database (Le Novère et al., 2006) as SBML (Systems Biology Markup Language) format (Hucka et al., 2003), with the identifier BIOMD0000000503 (Messiha et al., 2014; Smallbone et al., 2013). The kinetic model represented the glycolytic pathway from glucose down to ethanol as well as the pentose phosphate pathway, and only the glycolytic part was used in our simulation. Briefly, we first perturbed the original model (WT) to generate models of four mutant strains (mutant 01 to 04), and then 50 datasets including the amounts of metabolites, enzymes, and metabolic fluxes were generated for each of the models. The parameters of the WT model in enzyme 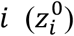 to be perturbed included kcat, Vmax, or an enzyme concentration. The magnitude of perturbation ζ_*l*_ (*l* = 1, … ,5), strain-specific noise, was set so that mutant strains with perturbed parameter sets ***z***^*l*^ were gradually deviated from the WT model (Table S8).

Based on the perturbed model, sample datasets were generated by introducing sample-specific noises 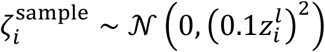 for each parameter value 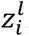, which represented variety between samples in the same strain and were assumed to be common in all the strains. Steady-state simulation using the parameter set with the strain-specific and sample-specific noises produced a dataset containing 50 samples with the amounts of metabolites, enzymes, and metabolic fluxes under each of the five conditions. All the steady-state simulation were executed using the Simbiology toolbox in MATLAB (The MathWorks, Inc., Natick, Massachusetts, United States of America).

To evaluate the performance of OMELET, only the dataset of the amounts of enzymes and metabolites, not including metabolic fluxes, were used as input. Since the amounts of transcripts were not available, we obtained the joint posterior distribution

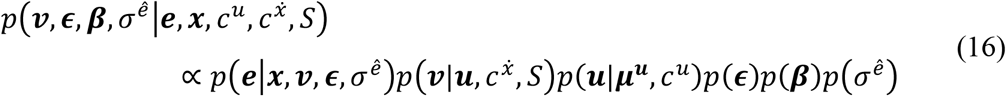

where the likelihood of transcripts (Table S6) was removed from equation (15). The independent fluxes were constrained so that the metabolic flux through glucose uptake (hexose transporter; HXT) in WT strain was fixed at one. A metabolic flux through each reaction was inferred as the relative value to that through HXT in WT. The parameters for coefficient of variances *c*^*u*^ and 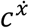 were fixed at 0.1 and 0.01, respectively. For MCMC sampling, we ran four chains of 20,000 iterations with 17,500 burnings with thinning 2, resulting in 5,000 samples in total. The metabolic fluxes inferred by OMELET were then compared with those obtained from the perturbation and steady-state simulation of the kinetic model.

### Contributions of regulators to changes in metabolic flux

We define a contribution *ψ*_*j*ℎ_ of regulator ℎ to changes in metabolic flux through reaction *j* between conditions. The regulators include transcripts, unaccounted enzyme regulators, substrates, products, cofactors, allosteric effectors, and unaccounted flux regulators. The unaccounted enzyme regulators can include other regulatory mechanisms of the protein amount of enzyme such as protein degradation and stability. The unaccounted flux regulators can include other regulators such as phosphorylation of enzymes and unknown allosteric effectors not included in OMELET. The concept of the contribution is to partition the cause of changes in metabolic flux between conditions into underlying changes in the amounts of regulators including enzymes and metabolites. The contribution was calculated based on propagation of uncertainty of regulators’ amounts to metabolic flux, and a similar approach was described in a previous study (Hackett et al., 2016). Note that we analyzed only the local effects of regulators on changes in metabolic flux and do not evaluate the effects on changes in metabolic flux in which the regulator was not directly participated.

Before we calculated the contribution, we define the amounts of unaccounted flux regulators and unaccounted enzyme regulators. Based on equation (9), the inferred metabolic flux through reaction *j* in condition *l* can be described as

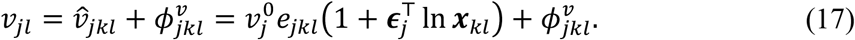

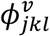 is the amount of an unaccounted flux regulator in sample *k* in condition *l* and represents the deviation of the inferred metabolic flux from that calculated using linlog kinetics. Similarly, based on equation (13), the measured amount of enzyme in reaction *j* can be described as

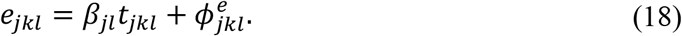

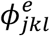 is the amount of an unaccounted enzyme regulator and represents the deviation of the measured amount of the enzyme from that calculated using the amount of transcript and the protein turnover coefficient. Combining equations (17) and (18), we obtain

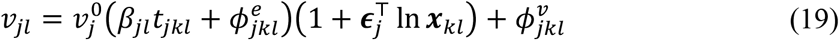

which represents metabolic flux *v*_*jl*_ as a function of the transcript *t*_*jkl*_, the unaccounted enzyme regulator 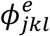, the metabolites ***x***_*kl*_ including substrates, products, cofactors and allosteric effectors, as well as the unaccounted flux regulator 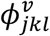.

We defined the contribution to changes in metabolic flux from transcripts, unaccounted enzyme regulators, and metabolites of substrates, products, cofactors, and allosteric effectors, as well as unaccounted flux regulators. Based on propagation of uncertainty, assuming that interactions between regulators is ignored, the variance Var(*v*_*j*_) of inferred metabolic flux *v*_*j*_ through reaction *j* can be approximated as

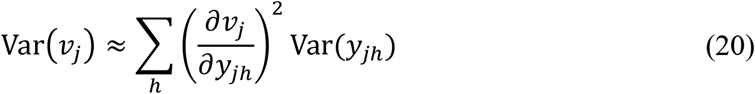

with the amount of regulator ℎ (*y*_*j*ℎ_), the sensitivity of the metabolic flux to the regulator ∂*v*_*j*_/∂*y*_*j*ℎ_, and the variance of the amount of the regulator Var(*y*_*j*ℎ_). Regulator ℎ is the transcript, the unaccounted enzyme regulator, substrate, product, cofactor, allosteric effector, or unaccounted flux regulator. The variance of the amount of the regulator between two conditions Var(*y*_*j*ℎ_) is expressed as the change in the amount of the regulator between the two conditions Δ*y*_*j*ℎ_ as

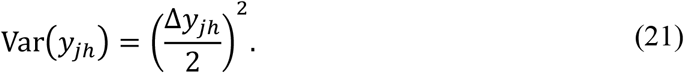

The sensitivity of the metabolic flux to each regulator ∂*v*_*j*_/∂*y*_*j*ℎ_ is defined based on equations (17) and (19) as

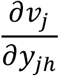

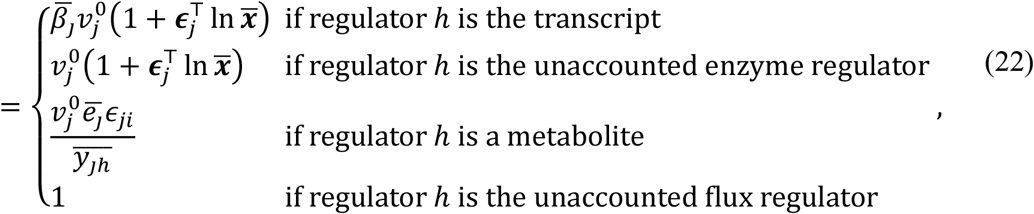

where 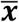 and 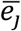 indicate the means of the metabolites and enzyme in reaction *j* across two conditions, respectively. Using the variance and the sensitivity of the metabolic flux to the amount of each regulator, we defined a contribution of the change in regulator ℎ to the change in metabolic flux through reaction *j* as

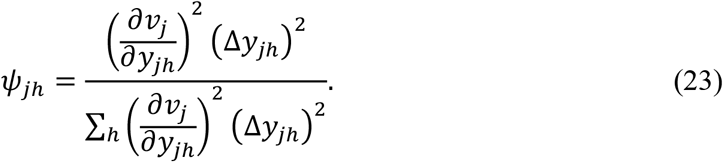

The contribution was a compositional data whose sum of the contributions of all the regulators to a change in metabolic flux equals one. The contribution ranged from zero to one, and the larger value meant the stronger effect of the regulator to the change in metabolic flux.

The contribution was calculated for changes in metabolic flux through each reaction between each pair of condition (Figure S7); WT and *ob*/*ob* mice in the fasting state, WT and *ob*/*ob* mice after oral glucose administration, fasting and after oral glucose administration in WT mice, and fasting and after oral glucose administration in *ob*/*ob* mice. Therefore, the calculated contributions represent the extent to which a change in each regulator contributed to the change in metabolic flux between the conditions.

### Metabolomic analysis

Metabolomic measurements were performed as previously described (Egami et al., 2021; Kokaji et al., 2020). Total metabolites and proteins were extracted from the liver with methanol:chloroform:water (2.5:2.5:1) extraction. Approximately 40 mg of the liver was suspended with 500 μL of ice-cold methanol containing internal standards [20 μM L-methionine sulfone (Wako), 2-morpholinoethanesulfonic acid (Dojindo), and D-Camphor-10-sulfonic acid (Wako)] for quantification of metabolites, then with 500 μL of chloroform, and finally with 200 μL of water. After centrifugation at 4,600 × g for 15 min at 4℃, the separated aqueous layer was filtered through a 5 kDa cutoff filter (Human Metabolome Technologies) to remove protein contamination. The filtrate (320 μL) was lyophilized and, prior to MS analysis, dissolved in 50 μL water containing reference compounds [200 μM each of trimesic acid (Wako) and 3-aminopyrrolidine (Sigma-Aldrich)]. Proteins were precipitated by addition of 800 μL of ice-cold methanol to the interphase and organic layers and centrifuged at 12,000 × g for 15 min at 4℃. The pellet was washed with 1 mL of ice-cold 80% (v/v) methanol and resuspended in 1 mL of sample buffer containing 1% SDS and 50 mM Tris-Cl pH8.8, followed by sonication. The total protein concentration was determined by bicinchoninic acid (BCA) assay and was used for normalization of metabolite concentration among samples.

All CE-MS experiments were performed using an Agilent 1600 Capillary Electrophoresis system (Agilent technologies), an Agilent 6230 TOF LC/MS system, an Agilent 1200 series isocratic pump, a G1603A Agilent CE-MS adapter kit, and a G1607A Agilent CE electrospray ionization (ESI)-MS sprayer kit. Briefly, to analyze cationic compounds, a fused silica capillary [50 μm internal diameter (i.d.) × 100 cm] was used with 1 M formic acid as the electrolyte (Soga and Heiger, 2000). Methanol/water (50% v/v) containing 0.01 μM hexakis(2,2-difluoroethoxy)phosphazene was delivered as the sheath liquid at 10 μL/min. ESI-TOFMS was performed in positive ion mode, and the capillary voltage was set to 4 kV. Automatic recalibration of each acquired spectrum was achieved using the masses of the reference standards ([^13^C isotopic ion of a protonated methanol dimer (2CH_3_OH+H)]^+^, *m*/*z* 66.0631) and ([hexakis(2,2-difluoroethoxy)phosphazene +H]^+^, *m*/*z* 622.0290). The metabolites were identified by comparing their m/z values and relative migration times to the metabolite standards. Quantification was performed by comparing peak areas to calibration curves generated using internal standardization techniques with methionine sulfone. The other conditions were identical to those described previously (Soga et al., 2006). To analyze anionic metabolites, a commercially available COSMO(+) (chemically coated with cationic polymer) capillary (50 μm i.d. × 105 cm) (Nacalai Tesque, Kyoto, Japan) was used with a 50 mM ammonium acetate solution (pH 8.5) as the electrolyte. Methanol/5 mM ammonium acetate (50% v/v) containing 0.01 μM hexakis(2,2-difluoroethoxy)phosphazene was delivered as the sheath liquid at 10 μL/min. ESI-TOFMS was performed in negative ion mode, and the capillary voltage was set to 3.5 kV. Automatic recalibration of each acquired spectrum was achieved using the masses of the reference standards ([^13^C isotopic ion of a deprotonated acetate dimer (2CH_3_COOH-H)]^−^, *m*/*z* 120.0384) and ([hexakis(2,2-difluoroethoxy)phosphazene +deprotonated acetate (CH_3_COOH-H)]^−^, *m*/*z* 680.0355). For anion analysis, D-camphor-10-sulfonic acid were used as the internal standards. The other conditions were identical to those described previously (Soga et al., 2009). The acquired raw data were analyzed using our proprietary software (Sugimoto et al., 2010).

### Proteomic analysis

#### Sample preparation of proteomic analysis

Sample preparation of proteome analysis were performed as described, previously (Matsumoto et al., 2017). Frozen powder of liver and muscle were lysed with a solution containing 2% SDS, 7 M urea, and 100 mM Tris-HCl, pH 8.8, and then subjected to ultrasonic treatment (five times for 30 s with intervals of 30 s) with a Bioruptor (Diagenode). The samples were diluted with an equal volume of water. The protein concentrations of the samples were determined with BCA assays (Bio-Rad), after which portions (200 μg of protein) were subjected to methanol–chloroform precipitation. The resulting pellet was dissolved in digestion buffer (0.5 M triethylammonium bicarbonate containing 7 M guanidine hydroxide) and heated at 56 °C for 30 min. Each sample was diluted with an equal volume of water, after which portions were subjected to BCA assays. The remaining solution (50 μl) was diluted with 50 μl of water and subjected to digestion with lysyl-endopeptidase (2 μg, Wako) for 4 h at 37 °C. After the addition of 100 μl of water, the samples were further digested with trypsin (2 μg, Thermo Fisher) for 14 h at 37 °C. To block cysteine/cystine residues, we treated the digest with 0.625 mM Tris(2-carboxyethyl)phosphine hydrochloride (Thermo Fisher) for 30 min at 37 °C, then performed alkylation with 3.125 mM 2-iodoacetoamide (Sigma) for 30 min at room tem perature and quenching with 2.5 mM N-acetyl-l-cysteine (Sigma). The resulting digests were freeze-dried and then labeled with the mTRAQΔ0 reagent (SCIEX). For a deep proteomics, tryptic digests were separated into 6 fractions with off-Line high-pH reverse phase chromatography (Matsumoto et al., 2017).

#### DDA of peptides for multiple reaction monitoring (MRM) method development

Target proteins were selected from the proteins listed in three Kyoto Encyclopedia of Genes and Genomes (KEGG) pathways; Glycolysis / Gluconeogenesis (mmu00010), Citrate cycle (mmu 00020), and Starch and sucrose metabolism (mmu 00500), and proteins related to insulin signaling. A key step for the establishment of a successful targeted proteomic analysis is the accurate selection of proteotypic peptides (PTPs) for the targets of interest. Therefore, we first performed a discovery phase aimed at the selection of PTPs, which was based on a deep proteomic characterization of total protein extract of the murine liver and muscle. mTRAQ-labeled peptides as mentioned above were fractionated by reversed-phase chromatography on a 16-cm column (inner diameter, 100 μm) packed in house with 2-μm L-column C18 material (CERI). Peptides were eluted with a linear gradient (typically 5–45% B for 40 min, 45–95% B for 1 min, and 95% B for 10 min, where A was 0.1% formic acid and 2% acetonitrile, and B was 0.1% formic acid and 90% acetonitrile), at a flow rate of 200 nl/min. The high-performance LC system (Eksigent nano-LC) was coupled to a TripleTOF5600 hybrid mass spectrometer (SCIEX). Data acquisition was performed in IDA mode with the iTRAQ option. Survey MS spectra were acquired for 100 ms, and the 10 most intense ions were isolated and then fragmented with an automatically optimized collision energy for an MS/MS acquisition time of 100 ms. Peak lists (mgf) generated by the AB SCIEX MS Data Converter were used to search a database containing IPI mouse version 3.44 (55 078 protein entries; IPI, European Bioinformatics Institute) protein sequences concatenated with decoy sequences, with the use of the MASCOT algorithm (Matrix Science). The search was conducted with the following parameter settings: trypsin was selected as the enzyme used, the allowed number of missed cleavages was set to two, and the mTRAQΔ0 label on the NH2-terminal or lysine residues and carbamidomethylation of cysteine were selected as fixed modifications. Oxidized methionine and the mTRAQΔ0 label on tyrosine were searched as variable modifications. The precursor mass tolerance was 50 p.p.m., and the tolerance of MS/MS ions was 0.02 mass/charge (m/z) units. We imported all significant peptide-spectrum matches (PSMs) (MASCOT score >20) into a relational database written in MySQL. From this dataset, we preferentially selected PTP candidates which met the following criteria: 1. more than six amino acids; 2. absence of tryptic missed-cleavage sites; 3. The C-terminus of PTPs is Lys or Arg; 4. absence of methionine residues. To verify whether these PTP candidates were actually identified and quantified in our MRM systems, trypsin digests used in “DDA of peptides for MRM method development” were labeled with either mTRAQΔ0 or mTRAQΔ4 and subjected to MRM assays. All MRM traces were analyzed by iMPAQT-quant (Matsumoto et al., 2017). Peak groups were scored on the basis of cosine similarity with MS/MS spectra obtained in DDA, peak coelution of at least three fragment ions for each peptide, the presence or absence of interfering ions, and intensity. Finally, we selected 2-3 PTPs per target protein and purchased from Funakoshi Co. All PTPs were resuspended with 20%-50% ethanol, pooled and labeled with mTRAQΔ4 using standard procedures.

#### MRM analysis

MRM analysis was performed with a QTRAP5500 instrument (SCIEX) equipped with nano-Advance UHPLC (MICHROM) and HTS-PAL/xt autosampler (CTC Analytics AG). Peptides were eluted with a linear gradient of 5%–30% B for 45 min, 30%–95% B for 46 min (where A is 0.1% formic acid and B is acetonitrile) at a flow rate of 200 nL/min. Parameters were set as follows: spray voltage, 2,000 V; curtain gas setting, 10; collision gas setting, high; ion-source gas-1 setting, 30 and interface-heater temperature, 150 °C. Collision energy (CE) was calculated with the following formulae: CE = (0.044 × m/z1) + 5.5 and CE = (0.051 × m/z1) + 0.5 (where m/z1 is the m/z of the precursor ion) for doubly and triply charged precursor ions, respectively. Collision-cell exit potential (CXP) was calculated according to the formula: CXP = (0.0391 × m/z2) − 2.2334 (where m/z2 is the m/z of the fragment ion). The declustering potential (DP) was set to 50, and the entrance potential (EP) was set to 10. Resolution for Q1 and Q3 was set to ‘unit’ (half-maximal peak width of 0.7 m/z). The scheduled MRM option was used for all data acquisition, with a target scan time of 2.0 s and MRM detection windows of 300s. More than three technical repeats were performed per sample. Raw data were analyzed by iMPAQT-Quant (Matsumoto et al., 2017) with the corresponding spectra library. Peak groups were scored on the basis of cosine similarity with the MS/MS spectra obtained in DDA, a peak co-elution of at least three fragment ions for each peptide, the presence or absence of interfering ions, and the intensity. Finally, all traces were manually checked to eliminate inadequate transitions. All quantified transitions were normalized across samples and converted into protein abundance by SRMstats software on R (Surinova et al., 2013).

### Transcriptomic analysis

Transcriptomic measurements were performed as previously described (Egami et al., 2021; Kokaji et al., 2020). Briefly, total RNA was extracted from the liver using RNeasy Mini Kit (QIAGEN) and QIAshredder (QIAGEN) and assessed for quantity using Nanodrop (Thermo Fisher Scientific) and for quality using the 2100 Bioanalyzer (Agilent Technologies). cDNA libraries were prepared using SureSelect strand-specific RNA library preparation kit (Agilent Technologies). The resulting cDNAs were subjected to 100-bp paired-end sequencing on an Illumina HiSeq2500 Platform (Illumina) (Matsumoto et al., 2007). Sequences were aligned to the mouse reference genome obtained from Ensembl database (Cunningham et al., 2015; Flicek et al., 2014) (GRCm38/mm10, Ensembl release 70) using the software package TopHat (Trapnell et al., 2009, 2012) (v.2.0.9), software in the Tuxedo tool. Cufflinks (v.2.2.1), software in the Tuxedo tool, was used to assemble transcript models from aligned sequences and to estimate the number of transcripts as an indicator of gene expression. The number of transcripts was shown as fragments per kilo base of exon per million mapped fragments.

### Blood glucose and insulin

The blood and insulin data at 0, 2, 5, 10, 15, 20, 30, 45, 60, 90, 120, 180, 240 minutes after oral glucose administration were previously reported (Kokaji et al., 2020).

## QUANTIFICATION AND STATISTICAL ANALYSIS

For the metabolites, enzymes, and transcripts, we defined increased and decreased molecules between the conditions using the following procedure (Figure 2; Table S2). For each molecule, we calculated the fold change of the mean amount of WT mice in the fasting state, WT mice after oral glucose administration, *ob*/*ob* mice in the fasting state, and *ob*/*ob* mice after oral glucose administration over the mean amount of WT mice in the fasting state. The significance of changes was tested by two-tailed Welch’s t-test for each molecule. The q values were calculated by Benjamini-Hochberg procedure. Molecules that showed an q value less than 0.05 are defined as significantly changed molecules. Among them, molecules with a fold change larger than 1.5 were defined as increased molecules between the conditions, whereas molecules with a fold change smaller than 0.67 were defined as decreased molecules. The Pearson correlation coefficient was calculated between the medians of metabolic fluxes inferred by OMELET and the means of metabolic fluxes simulated by the kinetic model (Figure S2C) or in the previous studies (Figures S5B and S5D) across all the reactions in all the conditions. The p-value was computed by transforming the correlation to create a t statistic having N-2 degree of freedom, where N is the number of samples.

## Notes

### Competing Interest Statement

The authors have declared no competing interest.

