## Supplemental Materials for "Multi-omics-based label-free metabolic flux inference reveals obesity-associated dysregulatory mechanisms in liver glucose metabolism"

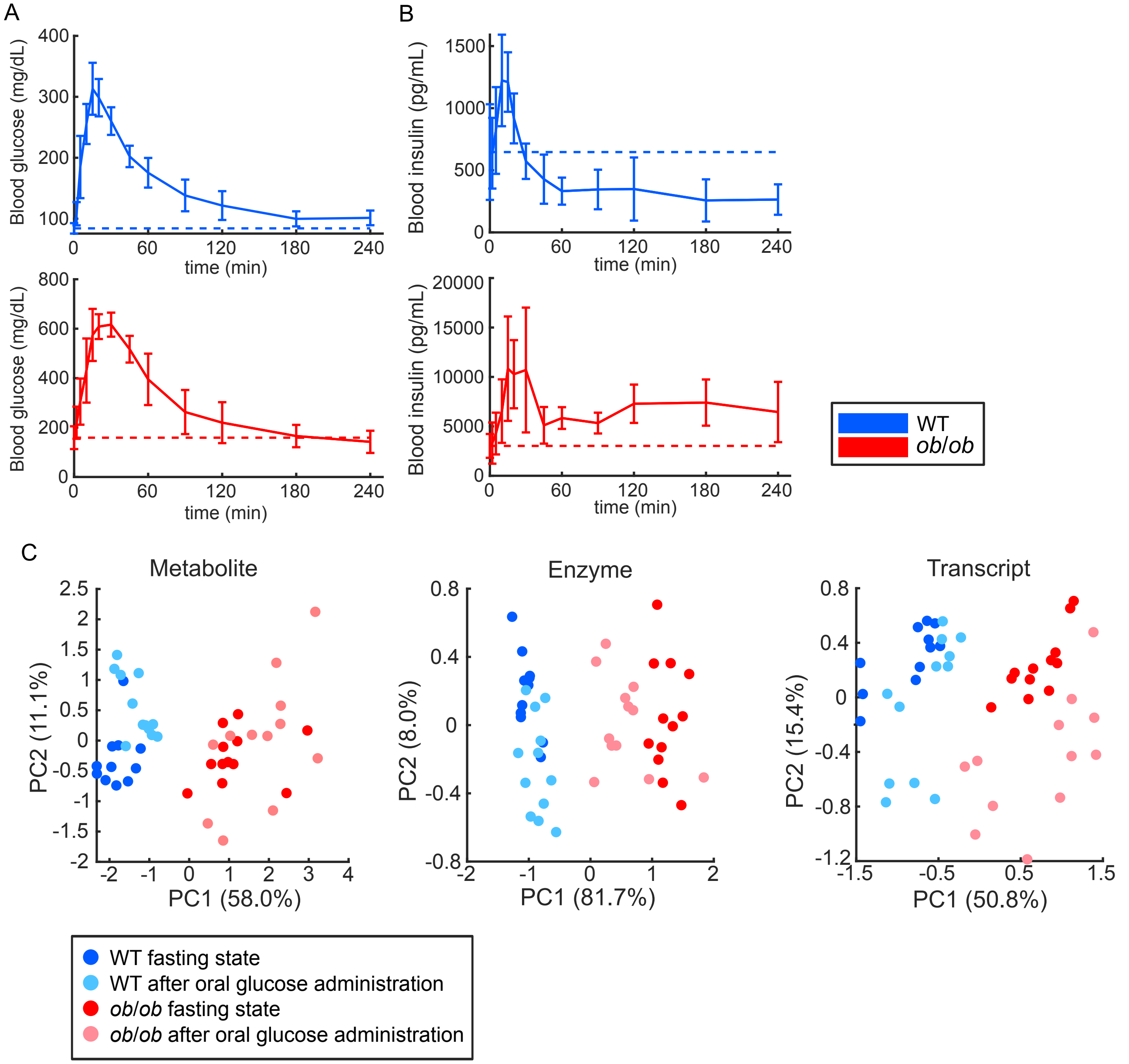


**Figure S1. Characterization of the metabolic states after oral glucose administration in WT and *ob*/*ob* mice. Related to Figure 2.**

(A, B) Blood glucose (A) and blood insulin (B) of WT (blue) and *ob*/*ob* mice (red) during oral glucose administration (Egami et al., 2021; Kokaji et al., 2020). The mean and SDs of five mice per genotype are shown. The dotted line shows the average value of each data at 0 min.

(C) Principal component analysis of the metabolites, enzymes, and transcripts for all the four conditions: WT in the fasting state (blue), WT after oral glucose administration (light blue), *ob*/*ob* in the fasting state (red), and *ob*/*ob* after oral glucose administration (pink). Score plots are shown for the first principal component (PC1) and the second principal component (PC2) with each dot of the same color representing an individual sample in the same condition. The contribution of the variance explained by each principal component to the total data variance (%) is shown in the x and y label.

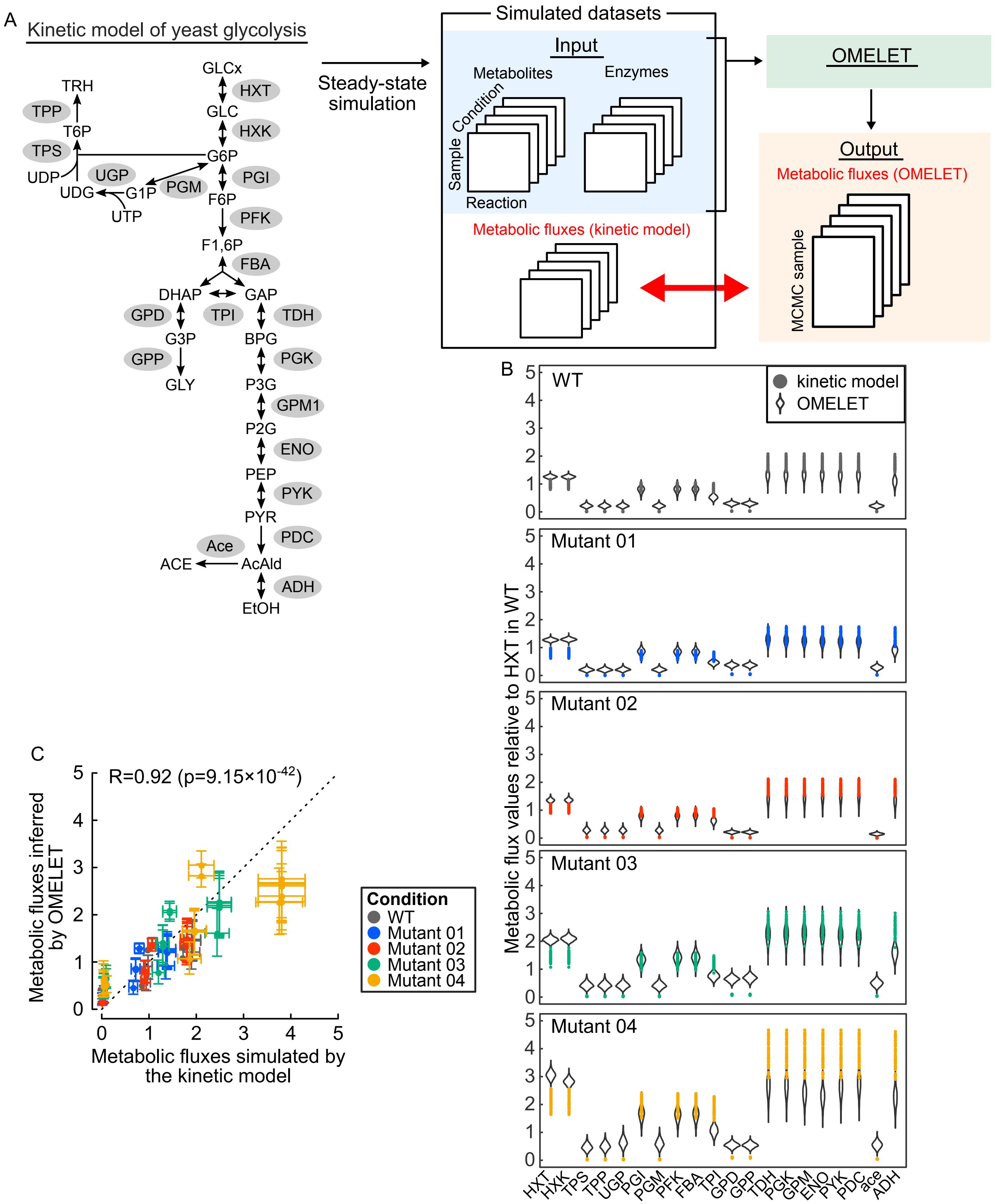
A
Kinetic model of yeast glycolysis
Steady-state simulation
Simulated datasets
Input
Metabolites
Enzymes
Condition
Sample
Reaction
Metabolic fluxes (kinetic model)
OMELET
Output
Metabolic fluxes (OMELET)
MCMC sample
GLCx
HXT
GLC
HXK
G6P
PGI
F6P
PFK
F1,6P
FBA
DHAP
GAP
TPI
GPD
TDH
G3P
BPG
GPP
PGK
GLY
P3G
GPM1
P2G
ENO
PEP
PYK
PYR
PDC
Ace
ACE
AcAld
ADH
EtOH
TRH
TPP
T6P
TPS
UDP
UGP
UDG
G1P
PGM
UTP
B
WT
kinetic model
OMELET
Mutant 01
Mutant 02
Mutant 03
Mutant 04
Metabolic flux values relative to HXT in WT
HXT HXK TPS TPP UGP PGI PGM PFK FBA TPI GPD GPP TDH PGK GPM ENO PYK PDC ace ADH
C
R=0.92 (p=9.15×10^-42)
Metabolic fluxes inferred by OMELET
Metabolic fluxes simulated by the kinetic model
Condition
WT
Mutant 01
Mutant 02
Mutant 03
Mutant 04

**Figure S2. Validation of the performance of OMELET using the kinetic model of yeast glycolysis. Related to Figure 3.**

(A) Overview of the validation of OMELET. The previously reported kinetic model representing the yeast glycolysis pathway consists of 23 metabolites and 21 reactions (Messiha et al., 2014; Smallbone et al., 2013). Cofactors and allosteric regulators are not presented on the map. We perturbed the original model (WT) to create models of four mutant strains (mutant 01 to 04), and the simulated datasets including metabolites, enzymes, and metabolic fluxes (n=50 per condition) were generated for each of the models. The magnitude of perturbation varied so that mutant strains gradually deviated from the WT model (Table S8; STAR Methods). To validate the performance of OMELET, only the enzyme and metabolite datasets, not including metabolic flux, were used as input of OMELET. The metabolic fluxes inferred by OMELET were compared with the metabolic fluxes generated by steady-state simulation of the kinetic model.

(B) Metabolic fluxes inferred by OMELET as posterior distribution (violin plots) and those simulated by the kinetic model (scatter plots) in each condition. All metabolic fluxes simulated by the kinetic model are normalized to the mean of that through hexose transport (HXT) in WT. The color represents the five conditions: WT (gray), mutant 01 (blue), mutant 02 (red), mutant 03 (green) and mutant 04 (yellow). See also Table S4.

(C) Correlations between the metabolic fluxes inferred by OMELET and those simulated by the kinetic model in each condition. For each condition, the mean ± SD of each metabolic flux simulated by the kinetic model (x-axis) is plotted against the median with 95% credible interval of the corresponding metabolic flux inferred by OMELET (y-axis). The Pearson correlation coefficient (R) and the p value between the metabolic fluxes simulated by the kinetic model and those inferred by OMELET are shown.

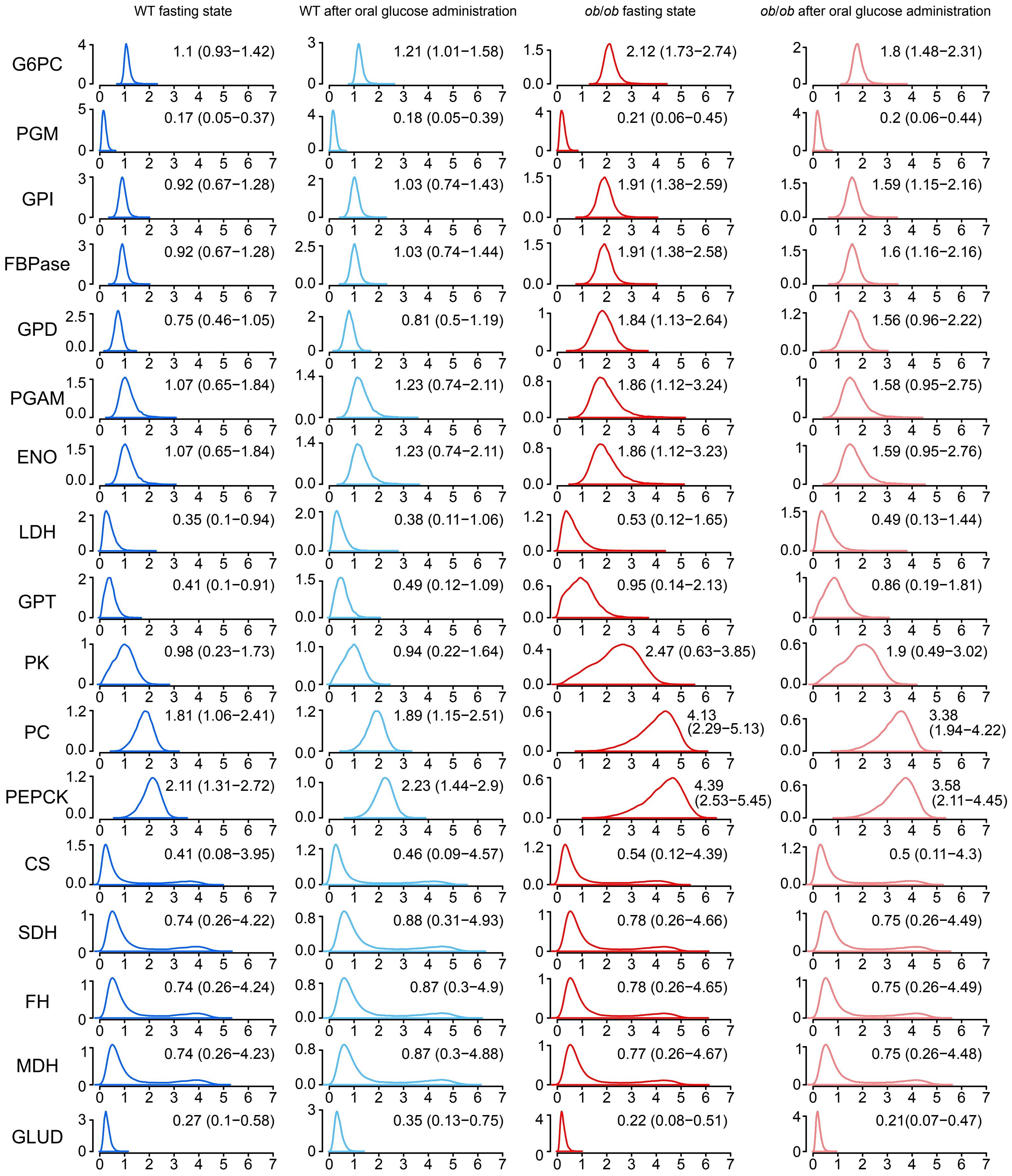
WT fasting state
WT after oral glucose administration
ob/ob fasting state
ob/ob after oral glucose administration
G6PC
1.1 (0.93−1.42)
1.21 (1.01−1.58)
2.12 (1.73−2.74)
1.8 (1.48−2.31)
PGM
0.17 (0.05−0.37)
0.18 (0.05−0.39)
0.21 (0.06−0.45)
0.2 (0.06−0.44)
GPI
0.92 (0.67−1.28)
1.03 (0.74−1.43)
1.91 (1.38−2.59)
1.59 (1.15−2.16)
FBPase
0.92 (0.67−1.28)
1.03 (0.74−1.44)
1.91 (1.38−2.58)
1.6 (1.16−2.16)
GPD
0.75 (0.46−1.05)
0.81 (0.5−1.19)
1.84 (1.13−2.64)
1.56 (0.96−2.22)
PGAM
1.07 (0.65−1.84)
1.23 (0.74−2.11)
1.86 (1.12−3.24)
1.58 (0.95−2.75)
ENO
1.07 (0.65−1.84)
1.23 (0.74−2.11)
1.86 (1.12−3.23)
1.59 (0.95−2.76)
LDH
0.35 (0.1−0.94)
0.38 (0.11−1.06)
0.53 (0.12−1.65)
0.49 (0.13−1.44)
GPT
0.41 (0.1−0.91)
0.49 (0.12−1.09)
0.95 (0.14−2.13)
0.86 (0.19−1.81)
PK
0.98 (0.23−1.73)
0.94 (0.22−1.64)
2.47 (0.63−3.85)
1.9 (0.49−3.02)
PC
1.81 (1.06−2.41)
1.89 (1.15−2.51)
4.13 (2.29−5.13)
3.38 (1.94−4.22)
PEPCK
2.11 (1.31−2.72)
2.23 (1.44−2.9)
4.39 (2.53−5.45)
3.58 (2.11−4.45)
CS
0.41 (0.08−3.95)
0.46 (0.09−4.57)
0.54 (0.12−4.39)
0.5 (0.11−4.3)
SDH
0.74 (0.26−4.22)
0.88 (0.31−4.93)
0.78 (0.26−4.66)
0.75 (0.26−4.49)
FH
0.74 (0.26−4.24)
0.87 (0.3−4.9)
0.78 (0.26−4.65)
0.75 (0.26−4.49)
MDH
0.74 (0.26−4.23)
0.87 (0.3−4.88)
0.77 (0.26−4.67)
0.75 (0.26−4.48)
GLUD
0.27 (0.1−0.58)
0.35 (0.13−0.75)
0.22 (0.08−0.51)
0.21(0.07−0.47)

**Figure S3. Posterior distributions of metabolic fluxes from OMELET. Related to Figure 4.**
Posterior distributions of the metabolic fluxes through all the analyzed reactions in the glucose metabolism. Each box contains a posterior distribution of a metabolic flux with its median and 95% credible interval.

Figure S4

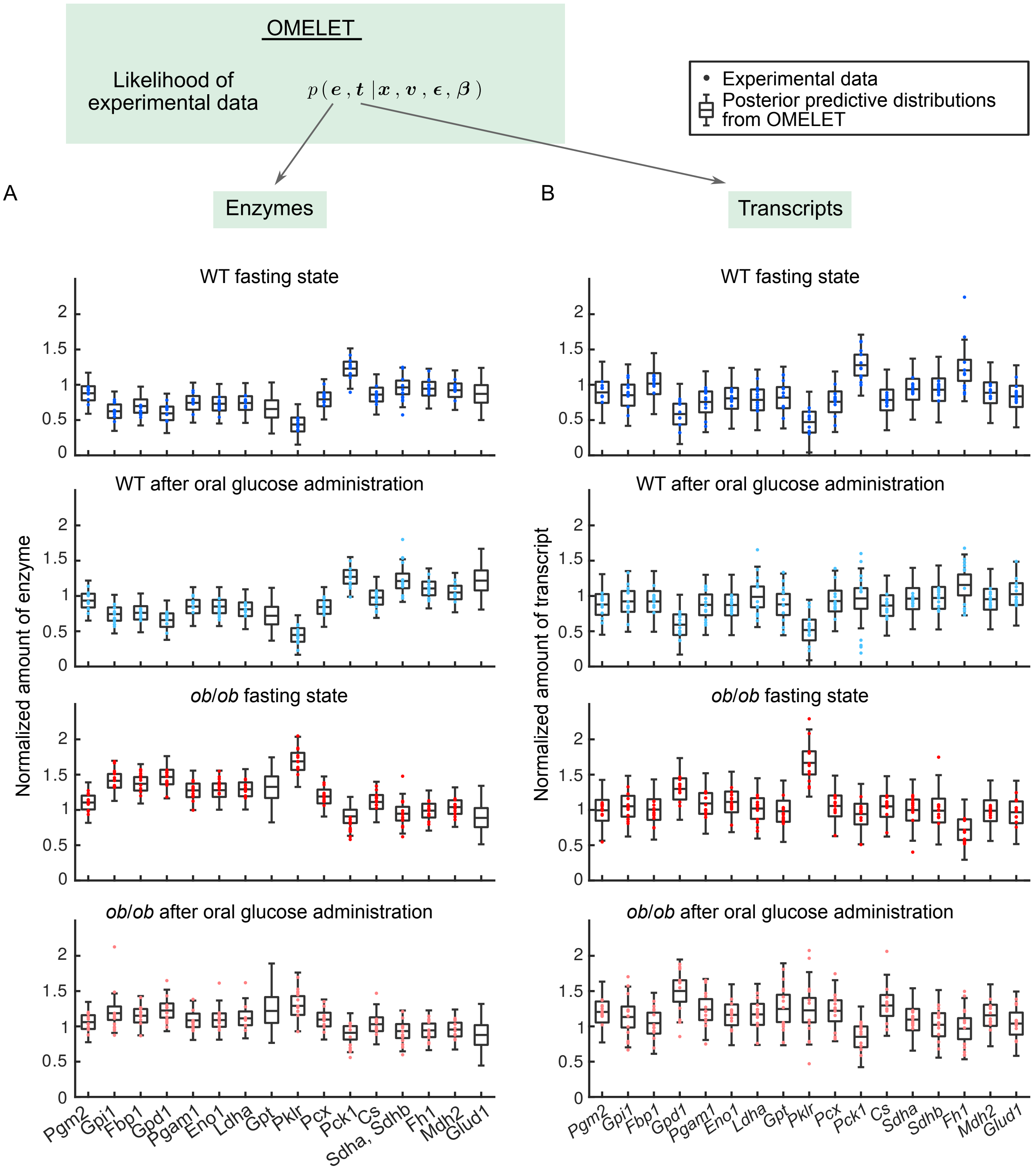

OMELET
Likelihood of experimental data
$p(\boldsymbol{e}, \boldsymbol{t} \mid \boldsymbol{x}, \boldsymbol{v}, \boldsymbol{\epsilon}, \boldsymbol{\beta})$
Experimental data
Posterior predictive distributions from OMELET
A
Enzymes
B
Transcripts
WT fasting state
WT after oral glucose administration
ob/ob fasting state
ob/ob after oral glucose administration
Normalized amount of enzyme
Normalized amount of transcript
Pgm2
Gpi1
Fbp1
Gpd1
Pgam1
Eno1
Ldha
Gpt
Pklr
Pcx
Pck1
Cs
Sdha, Sdhb
Sdha
Sdhb
Fh1
Mdh2
Glud1

**Figure S4. Fitting of the posterior predictive distributions from OMELET to the amounts of enzymes. Related to Figure 4.**

(A, B) Comparison between the amounts of enzymes (A) and transcripts (B) sampled from the posterior predictive distribution of the model for metabolic flux in OMELET (box plots) and those from the experimental data (scatter plots). The amounts of enzymes and transcripts are normalized to the mean of all the conditions. In Gpt and Glud1, the amounts of enzymes are not measured, and we plot only the posterior predictive distributions calculated from the estimated amounts of enzymes from transcripts in OMELET. The color represents four conditions: WT in the fasting state (blue), WT after oral glucose administration (light blue), *ob*/*ob* in the fasting state (red) and *ob*/*ob* after oral glucose administration (pink). The median of the posterior predictive distribution in the OMELET model is represented by a black line within the box for each reaction, the box extends from the lower to the $25^{th}$ and $75^{th}$ percentiles, and the whiskers extend to $2.5^{th}$ and $97.5^{th}$ percentiles to cover 95% of the data.

Figure S5

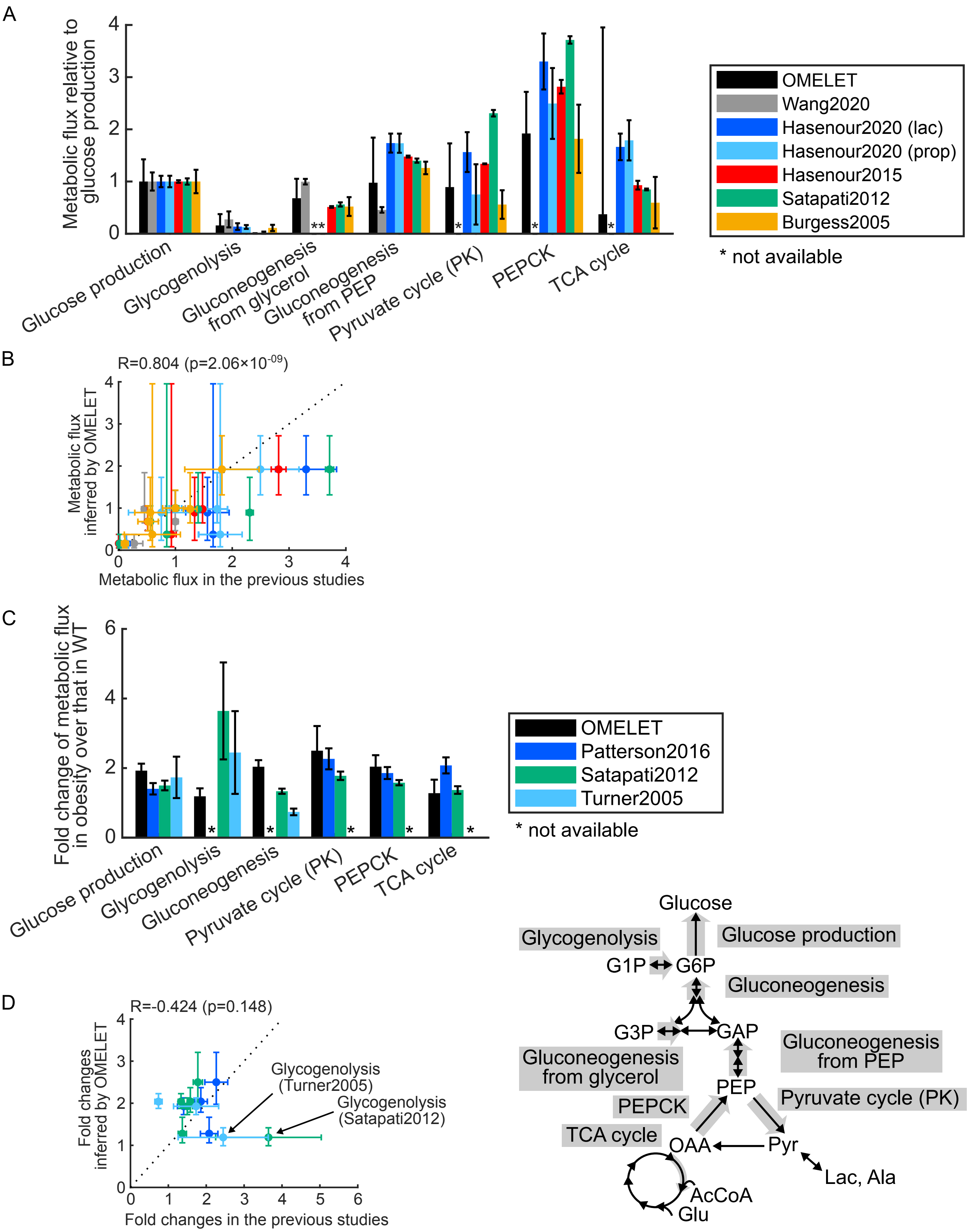
A
Metabolic flux relative to glucose production
OMELET
Wang2020
Hasenour2020 (lac)
Hasenour2020 (prop)
Hasenour2015
Satapati2012
Burgess2005
* not available
Glucose production
Glycogenolysis
Gluconeogenesis from glycerol
Gluconeogenesis from PEP
Pyruvate cycle (PK)
PEPCK
TCA cycle
B
R=0.804 (p=2.06×10^-09)
Metabolic flux inferred by OMELET
Metabolic flux in the previous studies
C
Fold change of metabolic flux in obesity over that in WT
OMELET
Patterson2016
Satapati2012
Turner2005
* not available
Glucose production
Glycogenolysis
Gluconeogenesis
Pyruvate cycle (PK)
PEPCK
TCA cycle
D
R=-0.424 (p=0.148)
Fold changes inferred by OMELET
Fold changes in the previous studies
Glycogenolysis (Turner2005)
Glycogenolysis (Satapati2012)
Glucose
Glycogenolysis
Glucose production
G1P
G6P
Gluconeogenesis
G3P
GAP
Gluconeogenesis from glycerol
Gluconeogenesis from PEP
PEP
PEPCK
Pyruvate cycle (PK)
TCA cycle
OAA
Pyr
Lac, Ala
AcCoA
Glu

**Figure S5. Validation of the metabolic fluxes inferred by OMELET with those in the previous studies. Related to Figure 4.**

(A) Metabolic fluxes inferred by OMELET with those in the previous metabolic flux analyses in fasting WT mice (Burgess et al., 2005; Hasenour et al., 2015, 2020; Satapati et al., 2012; Wang et al., 2020). The metabolic fluxes in the glucose metabolism in liver used for validation of the results from OMELET are shown in the bottom right. The bars and error bars represent the mean ± SD of metabolic fluxes normalized to the mean of the glucose production flux in each study. Metabolic flux not measured or available in each study is represented as an asterisk. The colors of the bars represent studies to measure metabolic fluxes. Hasenour2020 (lac) and Hasenour2020 (prop) represent the results using $^{13}C$ lactate and $^{13}C$ propionate as isotopic tracers, respectively, from the same study (Hasenour et al., 2020).

(B) Correlation between the metabolic fluxes inferred by OMELET with those in the previous metabolic flux analyses in fasting WT mice. The mean ± SD of each metabolic flux in the previous studies (x-axis) is plotted against the median with 95% credible interval of the corresponding metabolic flux inferred by OMELET (y-axis). The Pearson correlation coefficient (R) and the p value between the metabolic fluxes inferred by OMELET and those in the previous studies are shown.

(C) The fold changes of the metabolic flux of *ob*/*ob* mice over that of WT mice inferred by OMELET with those in the previous metabolic flux analyses using isotopic tracers in fasting *ob*/*ob* mice (Turner et al., 2005) and high-fat diet-induced obese mice (Patterson et al., 2016; Satapati et al., 2012). The metabolic fluxes in the glucose metabolism in liver used for validation of the results from OMELET are shown in the bottom right. The bars and error bars represent the mean ± SD of the fold changes of the metabolic flux of *ob*/*ob* mice over that of WT mice. Metabolic flux not measured or available in each study is represented as an asterisk. The fold changes of the metabolic fluxes through most reactions inferred by OMELET were consistent with those in the previous metabolic flux analyses in fasting *ob*/*ob* mice (Turner et al., 2005) and high-fat diet-induced obese mice (Patterson et al., 2016; Satapati et al., 2012) except for that through glycogenolysis. This discrepancy may be caused by the effects of isotopic tracers on glycogen metabolism and the different metabolic network including glycogen synthesis via glucose and gluconeogenic precursors.

(D) Correlation between the fold changes of the metabolic fluxes inferred by OMELET with those in the previous metabolic flux analyses in fasting WT mice. For each study, the mean ± SD of the fold changes of each metabolic flux in the previous study (x-axis) is plotted against the median with 95% credible interval of the corresponding metabolic flux inferred by OMELET (y-axis). The Pearson correlation coefficient (R) and the p value between the fold changes of the metabolic fluxes inferred by OMELET and those in the previous studies are shown. The correlation was not relatively high because of the discrepancy of the fold change of the metabolic flux through glycogenolysis.

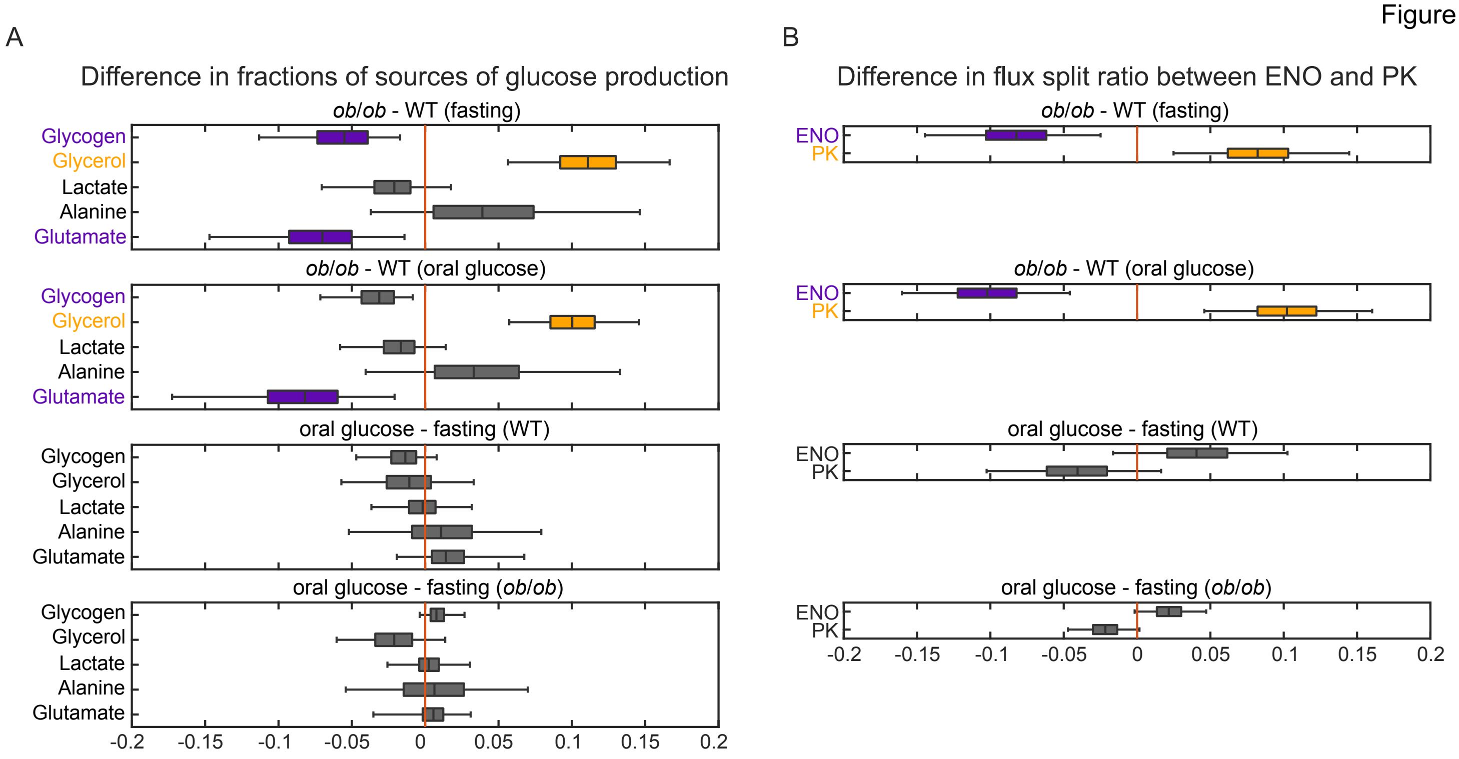


**Figure S6. Posterior distributions of differences in the fractions of sources of glucose production and those in the flux split ratios between conditions. Related to Figure 4.**

(A, B) The difference in the fractions of sources of glucose production (A) and that in the flux split ratio (B) between conditions. The median of the distribution of the difference is represented by a black line within the box for each fraction, the box extends from the lower to the 25th and 75th percentiles, and the whiskers extend to 2.5th and 97.5th percentiles to cover 95% of the data. The vertical orange line indicates the boundary where a difference equals zero. Of the boxes for each fraction or ratio, those with 95% credible intervals greater than zero are colored in yellow, and those with 95% credible intervals less than zero are colored in purple.

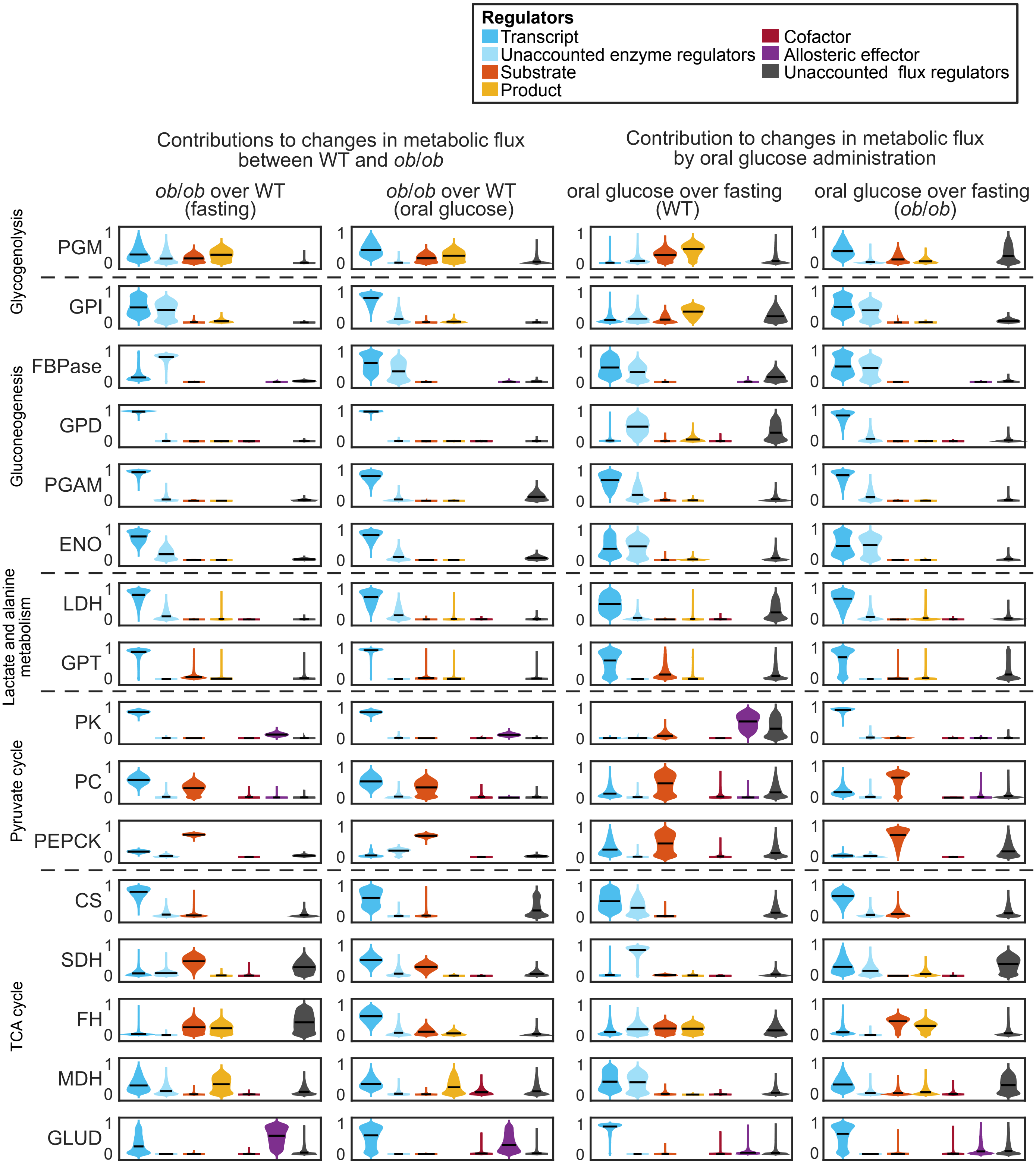


**Figure S7. Distributions of the contributions of regulators to changes in metabolic flux between conditions. Related to Figures 5 and 7.**

The contributions are calculated to changes in metabolic flux between WT and *ob*/*ob* mice in the fasting state and after oral glucose administration, as well as by oral glucose administration in WT and *ob*/*ob* mice. The means of the distributions are displayed as bar plots in Figures 5 and 7. The violin plots indicate the distribution of the contributions independently calculated in all the Markov chain Monte Carlo samples, and the horizontal black line in each violin plot represents the median.

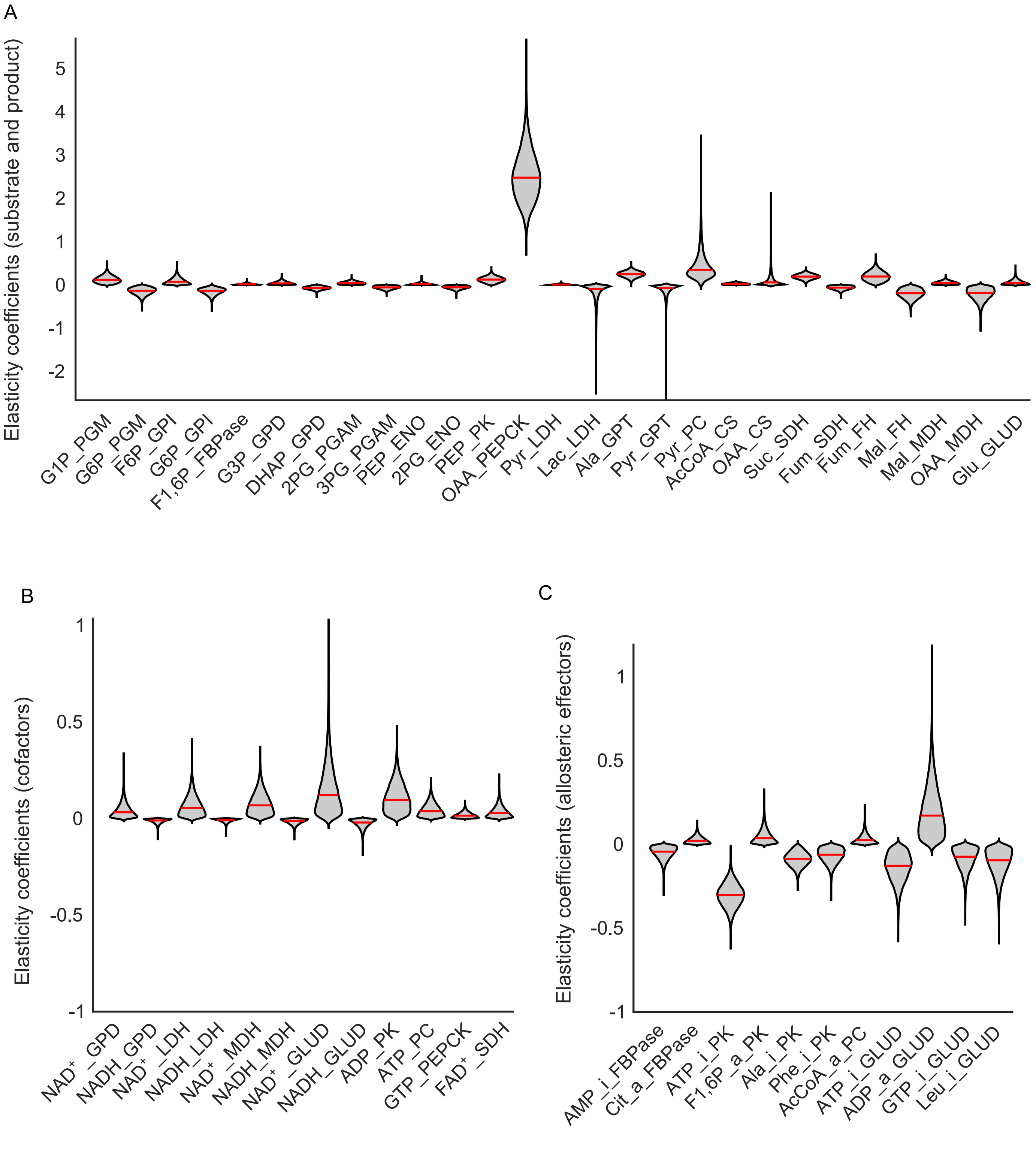


**Figure S8. Posterior distributions of elasticity coefficients from OMELET. Related to Figure 5.**
Posterior distributions of elasticity coefficients for substrates and products (A), cofactors (B), and allosteric effectors (C) in each reaction from OMELET. The horizontal red lines in each violin plot indicate the median. Labels of elasticity coefficients indicate names of metabolites with a name of the reaction that the metabolite is involved in. For example, "G1P_PGM" means the elasticity coefficient of G1P as substrate of reaction PGM. Allosteric inhibitors and activators are denoted as "_i_" and "_a_", respectively in (C).

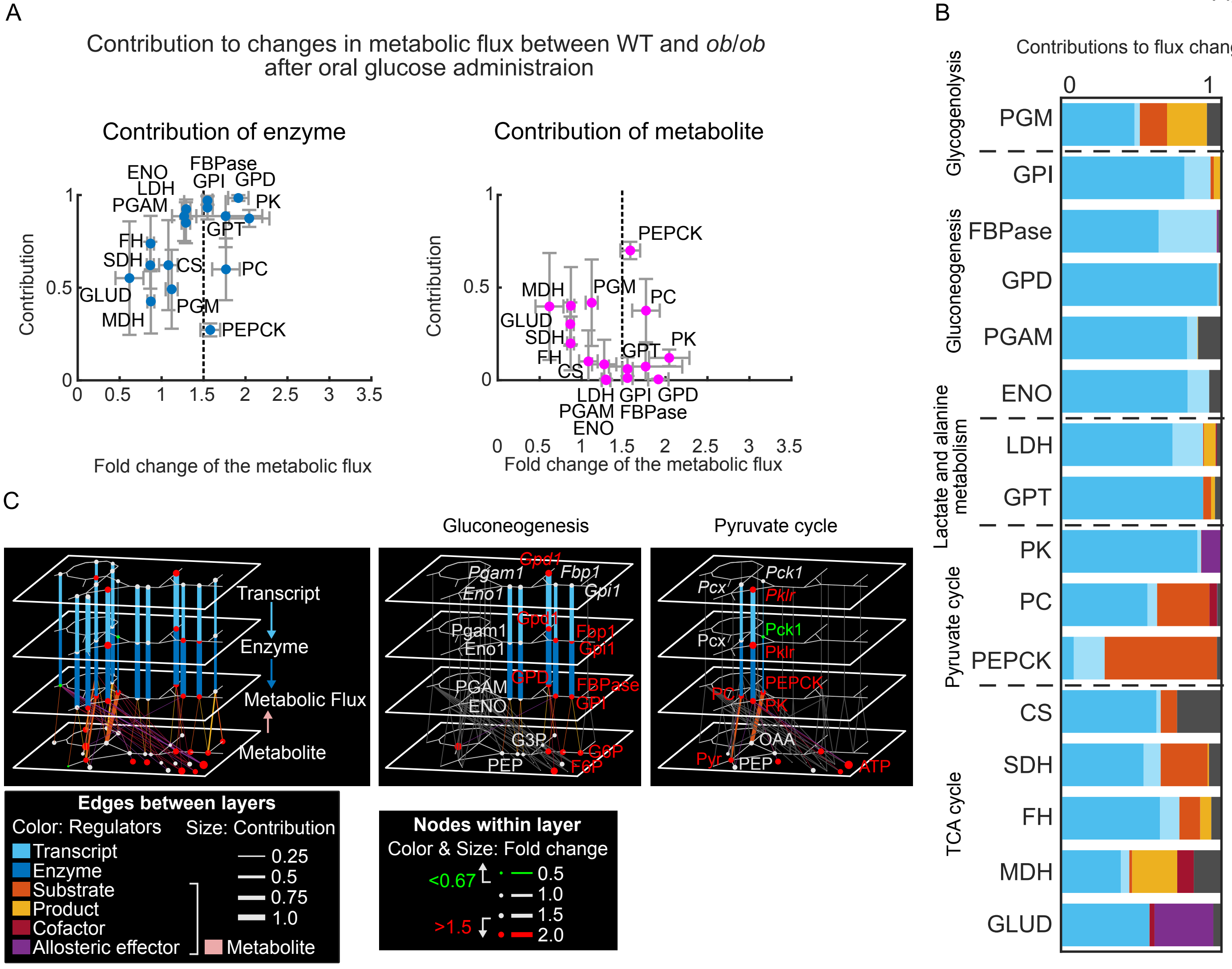


**Figure S9. Contributions of regulators to changes in metabolic flux between WT and *ob*/*ob* mice after oral glucose administration. Related to Figures 5 and 6.**

(A) Scatter plots illustrating the relationships between the contributions of enzyme (blue) and metabolite (pink) to changes in metabolic flux and the fold changes of the metabolic flux of *ob*/*ob* mice over that of WT mice after oral glucose administration. For each reaction, the mean ± SD of the distribution of the contributions of enzyme or metabolite to changes in metabolic flux (x-axis) is plotted against the mean ± SD of the distribution of the fold changes of the metabolic flux of *ob*/*ob* mice over that of WT mice after oral glucose administration (y-axis).

(B) Contribution of regulators to changes in metabolic flux between WT and *ob*/*ob* mice after oral glucose administration. The stacked bars indicate the mean of the contributions independently calculated in all the Markov chain Monte Carlo samples in Figure S7.

(C) Quantitative trans-omic networks for changes in metabolic flux between WT and *ob*/*ob* mice after oral glucose administration. The networks have the same structures as those in Figures 6.

**Table S1. Metabolites and reactions in the metabolic network for glucose metabolism in mice. Related to Figure 1 and STAR Methods.**

**Table S2. Metabolomic, proteomic, and transcriptomic data in the glucose metabolism in livers of WT and *ob*/*ob* mice in the fasting state and after oral glucose administration. Related to Figures 2, 6, and 7.**

**Table S3. Summary of differences in metabolites, enzymes, and transcripts among the different conditions. Related to Figure 2.**

**Table S4. Validation of OMELET using a kinetic model of yeast glycolysis. Related to Figure S2.**

**Table S5. Metabolic fluxes and kinetic parameters in the glucose metabolism inferred by OMELET. Related to Figures 4, 6, and 7.**

**Table S6. Contributions of regulators to changes in metabolic flux between conditions. Related to Figures 5-7.**

**Table S7. The priors and likelihoods of OMELET. Related to Figure 3.**

**Table S8. Perturbations to create strains using the kinetic model. Related to Figure S2.**

**Table S3. Summary of differences in metabolites, enzymes, and transcripts among the different conditions. Related to Figure 2.**

| | *ob*/*ob* over WT (fasting) | *ob*/*ob* over WT (oral glucose) | Oral glucose over fasting (WT) | Oral glucose over fasting (*ob*/*ob*) |
|---|---|---|---|---|
| Increased metabolites | Glycogen<br>G1P<br>G6P<br>F6P<br>F1,6P<br>PEP<br>Ala<br>Citrate<br>Succinate<br>ATP<br>ADP<br>NADH<br>GTP<br>Phe | Glycogen<br>G1P<br>G6P<br>F6P<br>F1,6P<br>Citrate<br>Succinate<br>ATP<br>ADP<br>AMP<br>NADH<br>Phe<br>Leu | Glycogen<br>G1P<br>G6P<br>F6P<br>PEP<br>ATP<br>GTP | None |
| Decreased metabolites | Lactate<br>Acetyl-CoA<br>$FAD^+$ | Lactate | None | None |
| Increased enzymes | Gpi1<br>Fbp1<br>Pgam1<br>Eno1<br>Gpd1<br>Pklr<br>Ldha | Gpi1<br>Fbp1<br>Gpd1<br>Pklr | None | None |
| Decreased enzymes | Pck1 | Pck1 | None | None |
| Increased transcripts | *Gpd1*<br>*Pklr* | *Gpd1*<br>*Pklr*<br>*Cs* | None | None |
| Decreased transcripts | *Fh1* | None | None | None |

**Table S7. The priors and likelihoods of OMELET. Related to Figures 3.**

| | |
|---|---|
| Prior for metabolic flux $\boldsymbol{v}$<br>- Independent flux $\boldsymbol{u}$<br>- Relaxation from the steady state $c^{\dot{x}}$<br>- Stoichiometric matrix $\boldsymbol{S}$ | $p(\boldsymbol{v}\mid\boldsymbol{u}) = \prod_{l=1}^{g} p(\boldsymbol{v}_l\mid\boldsymbol{u}_l)$<br>$p(\boldsymbol{v}_l\mid\boldsymbol{u}_l) = \mathcal{N}(\boldsymbol{v}_l\mid\boldsymbol{\mu}_l^{\boldsymbol{v}}, \boldsymbol{\Sigma}_l^{\boldsymbol{v}})$<br>$\boldsymbol{\mu}_l^{\boldsymbol{v}} = \boldsymbol{W}^{\boldsymbol{u}}\boldsymbol{\mu}_l^{\boldsymbol{u}}$<br>$\boldsymbol{\Sigma}_l^{\boldsymbol{v}} = \boldsymbol{W}^{\boldsymbol{u}}\boldsymbol{\Sigma}_l^{\boldsymbol{u}}(\boldsymbol{W}^{\boldsymbol{u}})^\top + \boldsymbol{W}^{\dot{\boldsymbol{x}}}\boldsymbol{\Sigma}_l^{\dot{\boldsymbol{x}}}(\boldsymbol{W}^{\dot{\boldsymbol{x}}})^\top$<br>Eq. (4)-(6) |
| Hyperprior for independent flux $\boldsymbol{u}$<br>- Mean of independent flux $\boldsymbol{\mu}^u$<br>- Variance of metabolic flux $c^u$ | $p(\boldsymbol{u}\mid\boldsymbol{\mu}^{\boldsymbol{u}}) = \prod_{l=1}^{g} p(\boldsymbol{u}_l\mid\boldsymbol{\mu}_l^{\boldsymbol{u}})$<br>$p(\boldsymbol{u}_l\mid\boldsymbol{\mu}_l^{\boldsymbol{u}}) = \mathcal{N}(\boldsymbol{u}_l\mid\boldsymbol{\mu}_l^{\boldsymbol{u}}, \boldsymbol{\Sigma}_l^{\boldsymbol{u}})$<br>Eq. (1)-(3) |
| Prior for elasticity coefficient $\boldsymbol{\epsilon}$ | $p(\boldsymbol{\epsilon}) = \prod_{j\in R}\prod_{i=1}^{m} p(\epsilon_{ji})$<br>$p(\epsilon_{ji}) = \begin{cases} \mathcal{H}(\epsilon_{ji}\mid 1) & \text{if metabolite } i \text{ is substrate or allosteric activator in reaction } j \\ -\mathcal{H}(\epsilon_{ji}\mid 1) & \text{if metabolite } i \text{ is product or allosteric inhibitor in reaction } j \\ \delta(\epsilon_{ji}\mid 0) & \text{others} \end{cases}$<br>Eq. (11) |
| Likelihood of enzymes<br>(Experimental data)<br>- Amounts of enzymes $\boldsymbol{e}$<br>- Amounts of metabolites $\boldsymbol{x}$<br>-<br>(Parameters)<br>- Metabolic flux $\boldsymbol{v}$<br>- Elasticity coefficient $\boldsymbol{\epsilon}$<br>- Error term $\sigma^{\hat{e}}$ | $p(\boldsymbol{e}\mid\boldsymbol{x},\boldsymbol{v},\boldsymbol{\epsilon},\sigma^{\hat{e}}) = \prod_{j\in R}\prod_{k=1}^{n_l}\prod_{l=1}^{g}\left[p(e_{jkl}\mid x_{jkl}, v_{jl}, \epsilon_{ji}, \sigma^{\hat{e}})\right]$<br>$p(e_{jkl}\mid\boldsymbol{x}_{kl}, v_{jl}, \boldsymbol{\epsilon}_j, \sigma^{\hat{e}}) = \mathcal{N}(e_{jkl}\mid\hat{e}_{jkl}, (\sigma^{\hat{e}})^2)$<br>$\hat{e}_{jkl} = \frac{v_{jl}}{v_j^0}\frac{1}{1+\boldsymbol{\epsilon}_j^\top \ln \boldsymbol{x}_{kl}}$<br>$v_j^0 = \frac{1}{g}\sum_{l=1}^{g}\mu_{jl}^{v}$<br>Eq. (9)-(10) |

| | |
|---|---|
| Prior for protein turnover coefficient $\boldsymbol{\beta}$<br><br>- Error term $\boldsymbol{\sigma}^{\beta}$ | $$p(\boldsymbol{\beta}) = \prod_{j \in R} \prod_{l=1}^{g} p(\beta_{jl})$$ $$p\left(\beta_{jl} \mid \sigma_l^{\beta}\right) = \mathcal{N}\left(\beta_{jl} \middle| 1, \left(\sigma_l^{\beta}\right)^2\right)$$ Eq. (14) |
| Likelihood of transcripts<br><br>(Experimental data)<br>- Amounts of transcripts $\boldsymbol{t}$<br><br>(Parameters)<br>- Protein turnover coefficient $\boldsymbol{\beta}$<br>- Error term $\sigma^{\hat{t}}$ | $$p\left(t \middle| \boldsymbol{x}, \boldsymbol{v}, \boldsymbol{\epsilon}, \boldsymbol{\beta}, \sigma^{\hat{t}}\right) = \prod_{j \in R} \prod_{k=1}^{n_l} \prod_{l=1}^{g} \left[p\left(t_{jkl} \middle| x_{jkl}, v_{jl}, \epsilon_{ji}, \beta_{jl}, \sigma^{\hat{t}}\right)\right]$$ $$p\left(t_{jkl} \middle| x_{jkl}, v_{jl}, \epsilon_{ji}, \beta_{jl}, \sigma^{\hat{t}}\right) = \mathcal{N}\left(t_{jkl} \mid \hat{t}_{jkl}, (\sigma^{\hat{t}})^2\right)$$ $$\hat{t}_{jkl} = \frac{1}{\beta_{jl}} \hat{e}_{jkl} = \frac{1}{\beta_{jl}} \frac{v_{jl}}{v_j^0} \frac{1}{1 + \boldsymbol{\epsilon}_j^{\top} \ln \boldsymbol{x}_{kl}}$$ Eq. (13) |

*Note*. $l = 1, \ldots, g$ represents a condition, $j \in R$ represents a reaction, $R$ represents a subset of the reactions to analyze, $i = 1, \ldots, m$ represents a metabolite, and $k = 1, \ldots, n_l$ represents a sample in each condition.

**Table S8. Perturbations to create strains using the kinetic model. Related to Figure S2.**

| Strain | Perturbation |
|---|---|
| Mutant 01 | $z_i^l = z_i^0 + \zeta_i^l,\ \zeta_i^l \sim \mathcal{N}(0, (0.4z_i)^2)$ |
| Mutant 02 | $z_i^l = z_i^0 + \zeta_i^l,\ \zeta_i^l \sim \mathcal{N}(0, (0.6z_i)^2)$ |
| Mutant 03 | $z_i^l = 2z_i^0 + \zeta_i^l,\ \zeta_i^l \sim \mathcal{N}(0, (0.4z_i)^2)$ |
| Mutant 04 | $z_i^l = 2z_i^0 + \zeta_i^l,\ \zeta_i^l \sim \mathcal{N}(0, (0.6z_i)^2)$ |
